# Clathrin’s adaptor interaction sites are repurposed to stabilize microtubules during mitosis

**DOI:** 10.1101/750034

**Authors:** Arnaud Rondelet, Yu-Chih Lin, Divya Singh, Arthur T. Porfetye, Harish C. Thakur, Andreas Hecker, Pia Brinkert, Nadine Schmidt, Shweta Bendre, Franziska Müller, Per O. Widlund, Tanja Bange, Michael Hiller, Ingrid R. Vetter, Alexander W. Bird

## Abstract

Clathrin plays an important role to ensure mitotic spindle stability and efficient chromosome alignment, independently of its well-characterized functions in vesicle trafficking. While clathrin clearly localizes to the mitotic spindle and kinetochore-fiber microtubule bundles, the mechanisms by which clathrin stabilizes microtubules have remained elusive. Here we show that clathrin adaptor interaction sites on clathrin heavy chain (CHC) are repurposed during mitosis to directly recruit the microtubule-stabilizing protein GTSE1 to the mitotic spindle. Structural analyses reveal that multiple clathrin-box motifs on GTSE1 interact directly with different clathrin adaptor interaction sites on CHC, in a manner structurally analogous to that which occurs between adaptor proteins and CHC near membranes. Specific disruption of this interaction in cells releases GTSE1 from spindles and causes defects in chromosome alignment. Surprisingly, this disruption causes destabilization of astral microtubules, but not kinetochore-microtubule attachments, and the resulting chromosome alignment defect is due to a failure of chromosome congression independent of kinetochore-microtubule attachment stability. Finally, we show that GTSE1 recruited to the spindle by clathrin stabilizes microtubules and promotes chromosome congression by inhibiting the activity of the microtubule depolymerase MCAK. This work thus uncovers a novel role of clathrin to stabilize non-kinetochore-fiber microtubules to support chromosome congression. This role is carried out via clathrin adaptor-type interactions of CHC with GTSE1, defining for the first time an important repurposing of this endocytic interaction mechanism during mitosis.

## INTRODUCTION

The precise and differential regulation of the stability of different populations of microtubules (MTs) during mitosis is critical for multiple aspects of cell division, including chromosome alignment and segregation, spindle positioning, and cytokinesis. The congression of chromosomes to the metaphase plate and their stable alignment is facilitated via multiple mechanisms which rely on astral MTs, kinetochore MTs (kMTs), and non-kMT inner-spindle MTs, as well as associated MT motor proteins including dynein, CENP-E, and chromokinesins (Maiato et al., 2017). Despite their critical importance, the basic mechanisms and regulation by which different MT populations are (de)stabilized over time and space to carry out these and other functions remain poorly understood.

Clathrin plays an integral role in mitotic MT organization/stabilization and chromosome alignment. During mitosis, clathrin localizes to the mitotic spindle and associates with kinetochore-fibers (k-fibers), bundles of 20-30 MTs in mammals that connect centrosomes to the kinetochores on chromosomes(Okamoto et al., 2000; Royle et al., 2005; Booth et al., 2011; McDonald et al., 1992; Nixon et al., 2015). Depletion of clathrin from cells leads to loss of MT stability in mitosis, fewer MTs within k-fibers, and defects in spindle integrity and alignment of chromosomes at the metaphase plate(Booth et al., 2011; Royle et al., 2005; Fu et al., 2010; Lin et al., 2010; Cheeseman et al., 2013). Importantly, these mitotic roles of clathrin are independent of its role in endocytosis and membrane trafficking(Royle et al., 2005; Smith and Chircop, 2012; Cheeseman et al., 2013; Royle, 2013). During mitosis, clathrin forms a complex with the proteins TACC3, PI3K-C2*α*, and the microtubule polymerase ch-Tog(Hubner et al., 2010; Lin et al., 2010; Fu et al., 2010; Booth et al., 2011; Gulluni et al., 2017). Formation of this complex (hereafter referred to as the CHC/TACC3 complex) and its recruitment to spindles depends on the direct interaction between clathrin heavy chain (CHC) and TACC3 phosphorylated on serine S558 by the Aurora A kinase, thereby restricting the function of this clathrin complex on microtubules to mitosis. (Hubner et al., 2010; Fu et al., 2010; Lin et al., 2010; Booth et al., 2011; Burgess et al., 2018; Hood et al., 2013; Burgess et al., 2015). While the recruitment of the CHC/TACC3 complex during mitosis to spindles is necessary for MT stabilization and chromosome alignment, the mechanisms by which it stabilizes MTs remains unclear.

Despite the initial characterization of TACC3 homologs in Drosophila (D-TACC) and Xenopus (XTACC3/Maskin) indicating a role in preferential stabilization of astral/centrosomal microtubules(Gergely et al., 2000; Barros, 2005; Kinoshita et al., 2005)most analyses of CHC/TACC3 complex function have focused on k-fibers, where several insights have come from electron microscopy (EM) analysis. Clathrin localizes near electron-dense “bridges” that have been observed connecting MTs within k-fibers(Hepler, 1970; Witt et al., 1981; Booth et al., 2011). A more recent EM tomography study has found that these “bridges” are more akin to a “mesh”, which interconnects MTs(Nixon et al., 2015). Depletion of TACC3 or clathrin from cells leads to a reduction in the number of “bridges”, as well as MTs, within a k-fiber(Booth et al., 2011). These and other observations have contributed to the hypothesis that CHC/TACC3 complexes may perform dual functions within k-fibers to organize and space MTs via physical bridges, as well as to stabilize MTs that makeup the k-fiber by lowering catastrophe rates(Booth et al., 2011; Royle, 2012). While a clathrin-based bridging mesh may organize k-fiber microtubule bundles, it remains unclear how clathrin stabilizes MTs.

Astral MT stability is governed not only by factors tracking the growing plus-ends of astral MTs (e.g. EB1, Kif18B) (Rogers et al., 2002; Stout et al., 2011; Tanenbaum et al., 2011), but also by proteins concentrated around centrosomes (e.g. gamma tubulin, pericentrin) (Zimmerman et al., 2004).TACC3 localizes to both MT plus-ends and spindle poles, and it was recently shown that TACC3 depletion in human and mouse cells, as previously shown in Drosophila and Xenopus, causes destabilization of interpolar and astral MTs (So et al., 2019; Singh et al., 2014; Gutierrez-Caballero et al., 2015; Nwagbara et al., 2014). Loss of centrosomal MT stability upon XTACC3 depletion has been shown to arise from increased activity of the microtubule depolymerase, MCAK/Kif2C, at centrosomes(Kinoshita et al., 2005). MCAK localizes to centromeres, growing MT plus-ends, and spindle poles during mitosis, where it controls astral/centrosomal microtuble stabilization (Lan et al., 2004; Kline-Smith et al., 2004) (Srayko et al., 2005; Rizk et al., 2009; Walczak et al., 1996; Wordeman and Mitchison, 1995) (Moore et al., 2005). It was proposed that a function of TACC3 is to antagonize MCAK’s potent activity at centrosomes to stabilize astral MTs, presumably by promoting activity of the microtubule depolymerase XMAP215/ch-Tog (Kinoshita et al., 2005). However, whether clathrin (or the CHC/TACC3 complex) is required for astral microtubule stabilization has not been addressed.

Another potential functional member of the CHC/TACC3 complex is GTSE1 (pronounced “*jitsee-one*” (Monte et al., 2000)), an intrinsically disordered protein that has been shown by proteomic analysis to interact with the CHC/TACC3 complex in cells(Hubner et al., 2010). Two recent studies showed that GTSE1 localizes to the spindle during mitosis, and like TACC3, is necessary for astral microtubule stabilization and efficient chromosome alignment(Bendre et al., 2016; Tipton et al., 2017). We showed that GTSE1 stabilizes MTs in mitosis by inhibiting MCAK activity (Bendre et al., 2016). In that study, cancer cells either depleted or knocked-out of GTSE1 showed destabilization of both astral MTs and kinetochore-MT attachments. Another study reported that a specific “slow turnover” population of MTs become stabilized upon depletion of GTSE1(Tipton et al., 2017). The reason for these seemingly contradictory results regarding MT stability after GTSE1 depletion is unclear. GTSE1 requires both CHC and TACC3 to localize to spindles, but does not appear to influence their localization (Hubner et al., 2010; Bendre et al., 2016; Cheeseman et al., 2013). The interactions and mechanism by which the CHC/TACC3 complex recruits GTSE1 to the spindle is unknown.

The ability of clathrin to function in endocytosis and vesicle trafficking is dependent on interactions between the CHC N-terminal domain (NTD) and clathrin “adaptor” proteins, named for their ability to provide a physical link between specific cargo receptors and clathrin to initiate recruitment of clathrin to membranes and ultimately form a lattice and coated vesicle(Lemmon and Traub, 2012; Robinson, 2015). The CHC NTD folds into a *β*-propeller across which four distinct adaptor interaction sites (“Site 1” through “Site 4”) are distributed. Site 1 of the NTD binds to adaptor proteins containing the “clathrin box motif” (CBM) consensus sequence LΦXΦ[DE] (where Φ refers to a hydrophobic residue and X to any amino acid(Dell’Angelica et al., 1998; Haar et al., 2000). Site 2 binds to the consensus sequence PWxxW, called the “W-box” (Miele et al., 2004). Site 3 has been reported to interact with the “arrestin box” sequence [LI][LI]GxL(Kang et al., 2009), but was recently shown to bind CBM motifs as well(Muenzner et al., 2016). Recently, a fourth site on the CHC NTD was identified (termed the “Royle Box”), although no consensus interacting motif has been elucidated(Willox and Royle, 2011; Muenzner et al., 2016). While it has been established that the NTD of CHC is necessary for its mitotic role, it is not required for CHC interaction with TACC3, and no adaptor-site interactions have been described whereby this domain is repurposed for a mitotic function(Royle et al., 2005; Royle and Lagnado, 2006; Hood et al., 2013; Smith et al., 2013; Lin et al., 2010). Likewise, despite a proteomic analysis of clathrin-dependent spindle proteins(Rao et al., 2016), there have been no reports of clathrin adaptor proteins localizing to the metaphase spindle and stabilizing MTs.

Here we show that adaptor interaction sites within the CHC NTD indeed play an important role in clathrin’s ability to stabilize MTs and promote chromosome congression during mitosis. These sites directly recruit GTSE1 to the spindle to inhibit MCAK activity and stabilize MTs. Therefore, in contrast to endocytic adaptor proteins, GTSE1 serves as a MT stability-promoting clathrin “effector”, whose function can be determined by clathrin localization. Abolishing the CHC-GTSE1 interaction in cells leads to defects in chromosome congression associated with a loss of centrosomal microtubule stability, but not kinetochore-MT attachment stability, implicating clathrin in stabilizing non-k-fiber MTs for chromosome congression.

## RESULTS

### Clathrin directly recruits GTSE1 to the spindle via its adaptor interaction sites

Previous mass spectrometry analysis indicated that CHC interacts with GTSE1 (Hubner 2010). We first confirmed this by showing that CHC and GTSE1 coimmunoprecipitate in mitotic cell lysates (Fig. 1 A). This interaction is direct, as purified GST-CHC abundantly pulled down purified GTSE1 (Fig. 1 B). To identify the interaction site of GTSE1 on CHC, we analyzed the binding of GTSE1 to fragments of GST-CHC. Only a construct containing the NTD of CHC (a.a.1-330) could pull down GTSE1 (Fig. 1 C). To further test whether CHC interacted with GTSE1 via established adaptor binding sites in the CHC NTD, we mutated Sites 1-3 individually in GST-CHC(1-330) and assayed for binding to GTSE1. Surprisingly, both Site 1 and Site 3, but not Site 2, were required for interaction with GTSE1 (Fig. 1 D).

**Figure 1:**
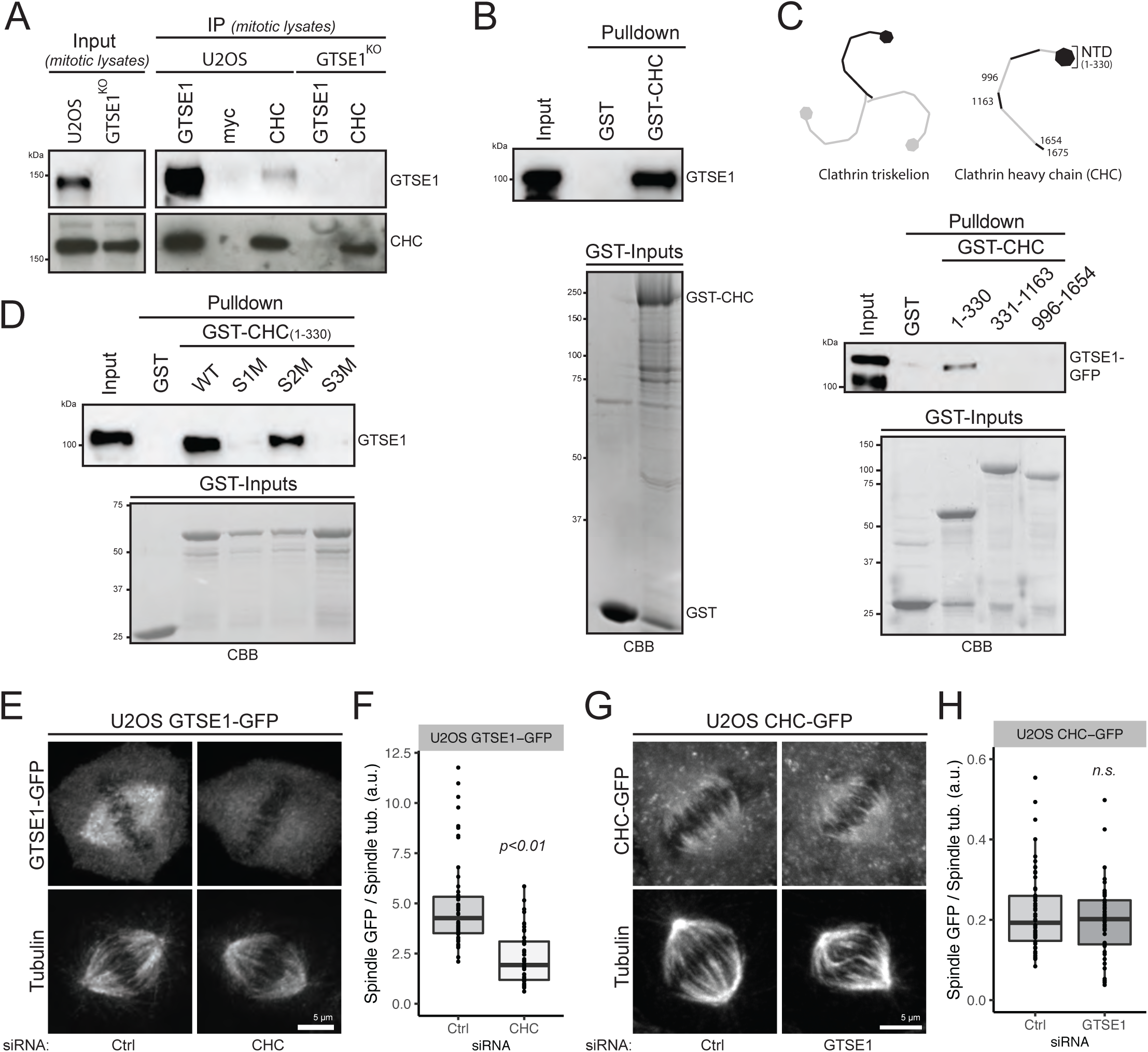
Adaptor interaction sites on CHC NTD directly interact with GTSE1 and recruit it to the spindle. **(A)** Immunoblot of mitotic cell lysates (input) and immunoprecipitations (IP) of GTSE1 or CHC (Myc antibody used as control). U2OS cells and U2OS cells knocked-out for GTSE1 (GTSE1^KO^) were arrested in mitosis with STLC. Immunoblots with anti-GTSE1 or anti-CHC antibodies. **(B)** GST-CHC pulldown of purified recombinant GTSE1. Immunoblot with anti-GTSE1 antibody. A Coomassie blue of GST inputs is presented. **(C)** Pulldown analysis of the interaction between purified GTSE1-GFP and CHC fragments fused to GST. Numbers indicate amino acids (a.a.) present in CHC fragments. Immunoblot with anti-GFP antibody. A Coomassie blue of GST inputs is presented. **(D)** Pulldown analysis of the interaction between purified GTSE1 and CHC(1-330) fused to GST. CHC was mutated at Site 1 (S1M: T87A, Q89A, K96E, K98E), Site 2 (S2M: Q152L, I154Q) or Site 3 (S3M: R188A, Q192A). Immunoblot with anti-GTSE1. A Coomassie blue of the GST inputs is presented. **(E)** Immunofluorescence images of clonal U2OS cells stably expressing a GTSE1-GFP BAC transgene (siRNA resistant) stained with antibodies against GFP and α-tubulin. Cells were transfected with Control (Ctrl) or CHC siRNA (66 h). All cells were concurrently transfected with GTSE1 siRNA to deplete endogenous GTSE1. **(F)** Quantification of the spindle recruitment of GTSE1-GFP in cells from (E). Mean GFP fluorescence intensity on half spindles was corrected for cytosolic background and normalized to the corresponding tubulin signal. (N=3 exp., at least 17 cells per condition and per experiment, 1 experiment presented, *p-value* from Wilcoxon test). **(G)** Immunofluorescence images of clonal U2OS cells stably expressing a CHC-GFP BAC transgene and transfected with Ctrl or GTSE1 siRNA. Cells were stained for α-tubulin, the GFP signal is from the fluorescing protein. **(H)** Quantification of CHC-GFP on the spindle of cells from (G). Mean GFP fluorescence intensity on half spindle was corrected for cytosolic background and normalized for tubulin intensity (N=3 exp., at least 22 cells per condition and per experiment, 1 experiment presented, *p-value* from Wilcoxon test). Scale bars: 5 µm. Numeric data in Table S1.

We next quantitatively analyzed the interdependency of CHC and GTSE1 for their recruitment to the mitotic spindle. RNAi-depletion of CHC caused GTSE1-GFP expressed from a bacterial artificial chromosome (BAC) transgene to delocalize from spindles (Fig. 1 E and F; Fig. S1 A and B). Conversely, CHC remained on spindles in U2OS cells after either RNAi-depletion or stable knockout of GTSE1 (Fig. 1 G and H; Fig. S1 C and D). Thus, while GTSE1 does not impact clathrin localization to spindles, clathrin recruits GTSE1 to spindles via a direct interaction with CHC adaptor interaction sites.

### GTSE1 binds to CHC in the manner of a clathrin adaptor protein

Because GTSE1 interacts with adaptor binding sites on CHC (Fig. 1 D), we searched for clathrin adaptor motifs in human GTSE1. This revealed five potential clathrin-binding motifs (further denoted motifs A, B, C, D and E), all located within the last 75 amino acids of GTSE1 (Fig. 2 A and B). Consistent with a role of these motifs in mediating the interaction between GTSE1 and CHC NTD, only GTSE1 fragments containing the terminal 75 amino acids could interact with CHC by pulldown assays (Fig. S2 A). To evaluate conservation of these potential motifs, we first performed sensitive sequence searches to identify GTSE1 homologs. This revealed that although a short domain present in the GTSE1 N-terminus can also be found within proteins in some but not all non-vertebrate species (Fig. S3, Interpro IPR032768), GTSE1 homologs containing the C-terminal domain that is required for clathrin binding are only conserved in vertebrates, with motifs C, D, and E in these homologs showing the highest conservation (Fig. 2 A; Fig. S3).

**Figure 2:**
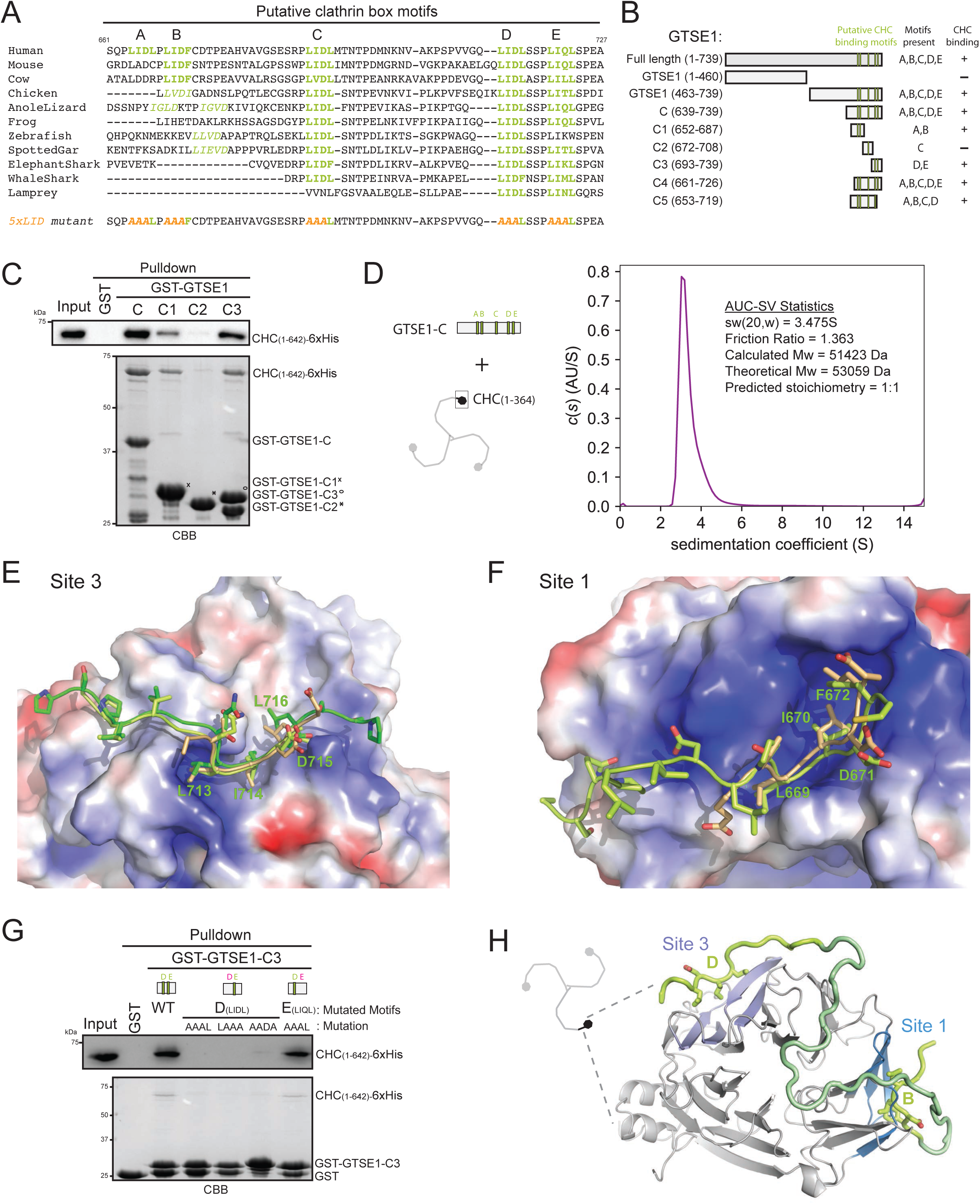
Multiple clathrin box motifs on GTSE1 and adaptor interaction sites on CHC are required for the GTSE1-CHC interaction. **(A)** Sequence alignment of GTSE1 C-terminus (a.a. 661 to 727 of human GTSE1). The 5 (A to E) putative clathrin-binding motifs are colored green and in bold. Divergent motifs are italicized. Mutations in the “5xLID” hGTSE1 mutant are shown in orange (last row). Alignment performed on full-length GTSE1 from indicated organisms using MUSCLE. **(B)** Scheme representing GTSE1 fragment constructs. Motifs A to E are shown. Numbers in brackets represent GTSE1 amino-acids. Binding of fragments to N-terminal CHC constructs is indicated. **(C)** Pulldown analysis of the interaction between GTSE1 fragments fused to GST (C, C1, C2 and C3) and CHC(1-642)-6xHis. Immunoblot (anti-His) and Coomassie blue are from separate experiments. (D) Analytical ultracentrifugation analysis of the stoichiometry of the complex between CHC(1-364) and GTSE1(639-739)-6xHis. The c(s) versus sedimentation coefficient plot with a statistic inset is presented. See also Fig. S2 E for raw data sedfit analysis curve. **(E)** Structures of GTSE1 motif D interacting with Site 3 on CHC. Superimposition (PyMOL) of the structures of CHC(1-364) in complex with GTSE1-C4 (dark green) (PDB ID: 6QNN) or with GTSE1-C5 (light green) (PDB ID: 6QNP). A reference structure (5M5S) of CHC bound to an amphiphysin peptide is presented in beige. Electrostatic potential on CHC are color coded (Blue: positive, Red: negative). **(F)** Structure of GTSE1 motif B (light green) interacting with CHC Site 1 (PDB ID: 6QNP). A reference structure (5M5S) of CHC bound to an amphiphysin peptide is presented in beige. See also Fig. S2 F and Table S2. **(G)** Pulldown analysis of the interaction between GST-C3 and CHC(1-642)-6xHis. C3 fragments were mutated at motif D or E and the corresponding mutated sequences are indicated. Immunoblot (anti-His) and Coomassie blue gels are from separate experiments. **(H) A** model of the interaction between GTSE1 and the CHC NTD with GTSE1 motifs B and D shown in Site 1 and Site 3 pockets, respectively. The GTSE1 C-terminus is colored in green. Light green regions represent structural data presented in (E) and (F) (PDB ID: 6QNP). Dark green sequences represent those absent from the electron density map in the crystal structure and were modeled manually with geometry minimization using Coot. CHC Site 1 is colored in blue, Site 3 in purple.

To determine the contribution of individual clathrin-binding motifs in GTSE1 to its interaction with the CHC NTD, we first performed a yeast two-hybrid analysis of the CHC-GTSE1 interaction after mutating the first three amino acids of each of the five motifs (A to E) to three alanines. While mutating any single motif lessened the interaction, mutation of motifs A, B or D had the most dramatic impact (Fig. S2 B). To confirm that multiple motifs on GTSE1 can directly interact with CHC, we tested whether short GTSE1 fragments containing either motifs A and B alone (GST-GTSE1-C1), motif C alone (GST-GTSE1-C2), or motifs D and E alone (GST-GTSE1-C3) (Fig. 2 B) could independently bind to CHC(1-642). Indeed, the fragment containing motifs A and B (C1), as well as the fragment containing motifs D and E (C3) (but not motif C alone (C2)), could efficiently pull-down CHC(1-642) (Fig. 2 C). Thus, multiple motifs in GTSE1 and multiple adaptor binding sites on CHC contribute to their interaction.

We next determined the molecular size of the complex formed between the CHC NTD and the GTSE1 C-terminus containing motifs A-E (fragment C) using analytical ultracentrifugation sedimentation velocity (AUC-SV) (Fig. 2 D; Fig. S2 C-E). This revealed a 51.4 kDa size of the complex, consistent with a heterodimer (theoretical Mw 53.0 kDa) and suggesting a 1:1 stoichiometry. Therefore, although there are multiple interaction interfaces between these CHC and GTSE1 proteins, they did not form multimeric complexes. Together these data suggest a model where multiple clathrin binding motifs on one GTSE1 molecule bind to multiple adaptor sites on one CHC NTD.

To understand how the complement of clathrin adaptor motifs on GTSE1 and adaptor binding sites on CHC interact, we sought to co-crystallize these regions of the proteins. We determined separate X-ray crystal structures for the complexes formed between the CHC NTD(1-364) and two different GTSE1 C-terminal fragments encompassing either motifs A-E or A-D (GTSE1-C4 and GTSE1-C5, respectively, Fig. 2 B and Table S2A). In both crystal structures, Sites 1 and 3, but not Site 2 or Site 4 on the CHC NTD, were consistently occupied by clear electron densities (Fig. 2, E and F; Fig. S2 F; Table S2B), consistent with our pull-down interaction data (Fig. 1 D). In both structures, the electron density present in Site 3 could unambiguously be attributed to the region encompassing GTSE1 motif D (Fig. 2 E; Fig. S2 F; Table S2B). Noteworthy, the GTSE1 residues (L713, I714 and L716) buried in the hydrophobic pocket of site 3 are those of a conserved clathrin box motif. Consistent with an integral role of motif D in binding to CHC, mutation of motif D, but not of motif E, on a GST-GTSE1-C3 fragment abolished its binding to CHC(1-642) (Fig. 2 G). In contrast to what we observed for Site 3, Site 1 was found occupied by three different GTSE1 motifs in the two crystals: motif E in the crystal with GTSE1-C4, and two different motifs, B and C, in different symmetry-related CHC molecules in the crystal with GTSE1-C5 (Fig. 2 F; Table S2B; for detailed interpretations and discussion of structural data see Supplementary Information). In all cases, bulky hydrophobic residues (i.e. L669, I670 and F672 from motif B; L689, I690 and L692 from motif C; L720, I721 and L723 from motif E) were buried in the hydrophobic cleft formed by blade 1 and 2 of the CHC NTD beta propeller, consistent with previously described structures of CBMs interacting with Site 1 (Fig. 2 F; Fig. S2 F; amphiphysin peptide bound to CHC is superimposed for comparison) (Muenzner et al., 2016) (Haar et al., 2000; Dell’Angelica et al., 1998). Mutational analyses indicated a central contribution of motif B in the interaction of GTSE1 with CHC, while motifs C and E contributed the least (Fig S2 B). Furthermore, the observed motif E interaction with Site 1 is likely a crystallization artefact (Fig. S2 B; see Supp Information). All together, these data show that the GTSE1-CHC interaction is reminiscent of the interaction between clathrin and its adaptor proteins in interphase, as clathrin box motifs in GTSE1 bind directly to adaptor binding Sites 1 and 3 on the CHC NTD. Multiple interaction sites between the two proteins appear to cooperate to facilitate maximum binding of a 1:1 complex, as schematically modeled in Fig. 2H.

### GTSE1 recruitment by the CHC/TACC3 complex is required for efficient chromosome alignment

To determine the importance of the CHC-GTSE1 interaction in mitosis, we generated two stable and clonal U2OS transgenic cell lines expressing either RNAi-resistant wild type GTSE1-GFP (GTSE1^WT^ cells) or GTSE1-GFP mutated at all 5 clathrin box motifs (A, B, C, D and E; further denoted “GTSE1^5xLID^“; see Fig. 2 A) from BACs at levels equivalent to endogenous GTSE1 (Fig. 3 A). We first confirmed that GTSE1^5xLID^-GFP was no longer able to interact with CHC in cells by immunoprecipitation followed by quantitative mass spectrometry and western blot (Fig. 3, B and C; Fig. S4 A; Table S3). We next asked whether loss of interaction with CHC abolished the ability of GTSE1 to associate with CHC/TACC3 complex members. Indeed, while GTSE1-GFP significantly associated with all known members of the CHC/TACC3 complex (CHC, TACC3, ch-TOG, and PI3K-C2a), GTSE1^5xLID^-GFP did not associate with any (Fig. 3, B and C; Fig. S4 A). Conversely, the ability to bind to MCAK, a known direct interactor of GTSE1, was preserved(Bendre et al., 2016)(Fig. 3, B and C, Table S3). This indicated that GTSE1 associates with the CHC/TACC3 complex exclusively via adaptor-interaction sites on CHC. Consistently, disruption of the CHC-TACC3 interaction by Aurora A inhibition prevented the association of GTSE1-GFP with TACC3 but not with CHC (Fig. S4 A). Likewise, we could not detect a direct interaction between purified GST-GTSE1 and pTACC3, even after mitotic kinase phosphorylation of GTSE1 (Fig. S4, B and C).

**Figure 3:**
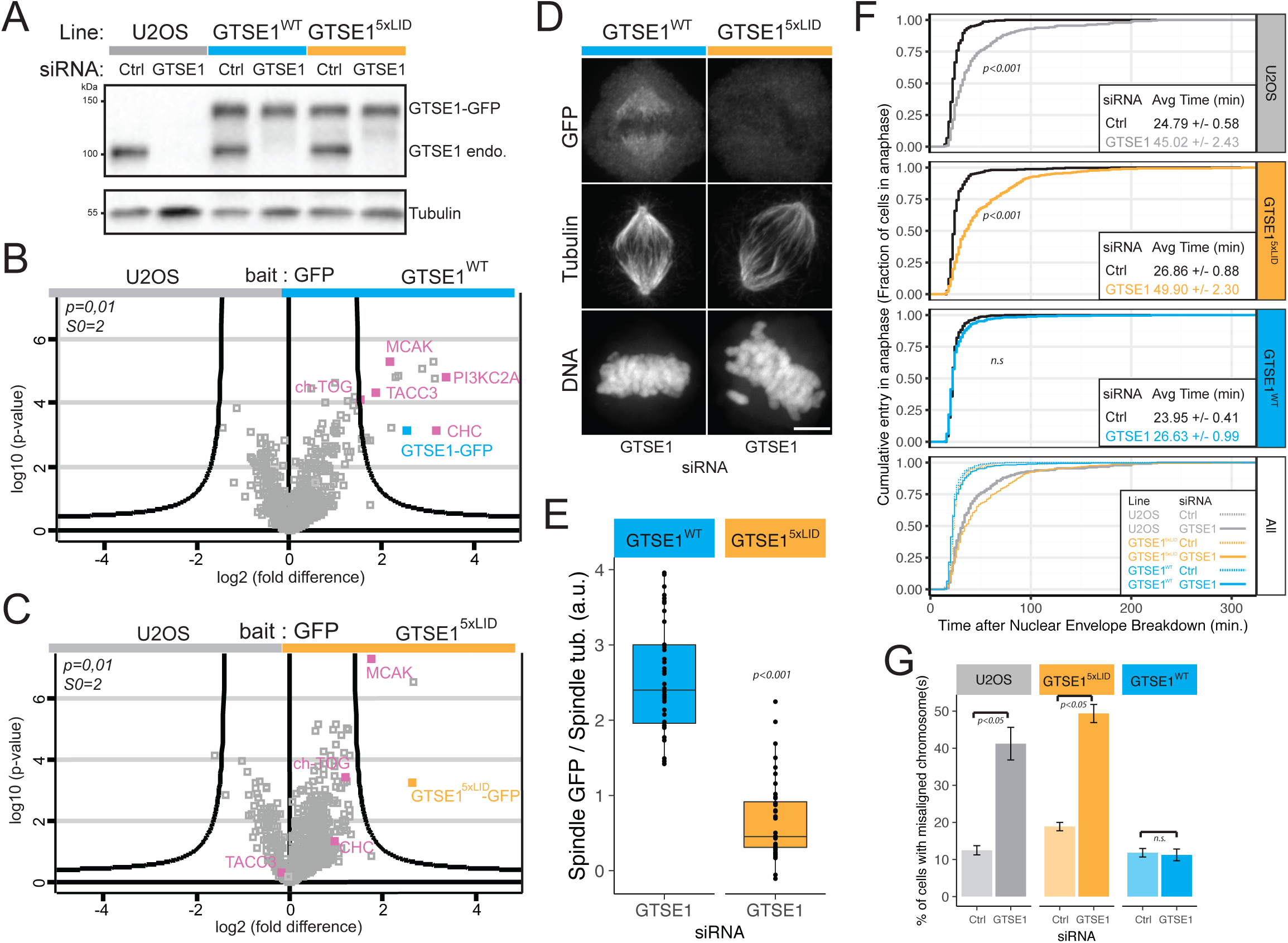
The CHC/TACC3 complex recruits GTSE1 to the spindle and promotes chromosome congression. **(A)** GTSE1^WT^ and GTSE1^5xLID^ cell lines express RNAi-resistant transgenes at endogenous levels. Western blots of U2OS clonal lines expressing RNAi-resistant GTSE1-GFP (GTSE1^WT^ cells) or GTSE1^5xLID^-GFP (Fig. 2 B, GTSE1^5xLID^ cells) from BAC transgenes. Cells were transfected with control (Ctrl) or GTSE1 siRNA, and cell lysates probed with antibodies against GTSE1 (recognizing both endogenous (“endo.”) and GFP-tagged (“GFP trans.”) GTSE1) and α-tubulin. **(B, C)** Disruption of the CHC-GTSE1 interaction prevents the association of GTSE1 with the CHC/TACC3 complex. Volcano plots show GTSE1-interacting proteins (pink) following anti-GFP immunoprecipitation of mitotic lysates from GTSE1^WT^, GTSE1^5xLID^, and U2OS cells, analyzed by mass spectrometry in triplicate. GTSE1-GFP is labeled in blue and GTSE1^5xLID^-GFP in orange. Curved lines represent significance threshold (t-test with threshold at *p=0.01*, S0=2, Table S3). **(D)** Immunofluorescence images of GTSE1^WT^ and GTSE1^5xLID^ cells transfected with GTSE1 siRNA. α-tubulin, GFP and DNA staining are shown. **(E)** Quantification of the GTSE1-GFP transgene on spindles of GTSE1^WT^ and GTSE1^5xLID^ cells depleted for endogenous GTSE1. Mean GFP fluorescence intensity on half spindles was corrected for cytosolic background and normalized for tubulin signal (n=58 cells per condition over N=3 exp., min. 13 cells per condition and per experiment, 1 experiment presented, *p-value* from Wilcoxon test). **(F)** Fraction of cells that have entered anaphase as a function of the time (min) after Nuclear Envelope Breakdown (NEB). The mean (+/-SE) time between NEB and anaphase onset is indicated in the insets. U2OS, GTSE1^WT^ and GTSE1^5xLID^ cells were transfected with Ctrl or GTSE1 siRNA. (n>223 cells per condition, N=3 over 4 exp., data were pooled for analysis and representation, *p-values* from Wilcoxon test show differences to respective Ctrl siRNA). **(G)** Percentage of fixed metaphase-like cells showing chromosome misalignment. The mean percentage (+/-SE) over N=3 experiments is presented (*p-values* from two-sided, unpaired t-test). Scale bars: 5 µm. Numeric data in Table S1.

We next analyzed the importance of the CHC-GTSE1 interaction for the spindle localization of GTSE1 and the CHC/TACC3 complex. GTSE1^5xLID^-GFPwas strongly impaired in its localization to the spindle by both fixed and live cell analysis (Fig. 3, D and E; Fig. S4 D). In contrast, and consistent with GTSE1 depletion data (Fig. 1, G and H; Fig. S1 D), recruitment of the CHC/TACC3 complex to the spindle was not changed in GTSE1^5xLID^ mutant cells, as TACC3 remained associated with the spindle (Fig. S4, E and F).

Depletion of GTSE1 leads to a mitotic delay linked to inefficient chromosome alignment(Bendre et al., 2016; Tipton et al., 2017). To test for a role of the CHC-GTSE1 interaction in chromosome alignment, we assayed both the time cells needed to transition from nuclear envelope breakdown (NEB) to anaphase onset (Fig. 3 F) and the percentage of metaphase cells showing misaligned chromosomes (Fig. 3 G). While the presence of a wildtype RNAi-resistant GTSE1-GFP transgene rescued the mitotic delay due to chromosome alignment defects in cells depleted for GTSE1, cells expressing only the GTSE1^5xLID^-GFP transgene showed significant defects in both mitotic duration and chromosome alignment. Together these data indicated that the CHC-GTSE1 interaction is required to recruit GTSE1 to spindles and facilitate chromosome alignment.

### CHC interaction with GTSE1 is required for the stabilization of non-kinetochore microtubules

We next probed for the mechanism by which the CHC-GTSE1 interaction promotes chromosome alignment. CHC is required for the stabilization and organization of MTs within the spindle(Lin et al., 2010; Royle et al., 2005; Booth et al., 2011; Cheeseman et al., 2013), and we previously showed that GTSE1 stabilizes multiple spindle MT subpopulations, including astral MTs and kMTs(Bendre et al., 2016). Spindles in GTSE1^5xLID^ cells displayed obvious morphological anomalies and often appeared “wavy”, with half spindles often displaying a “concave” rather than the normal “convex” shape, and a generally more prolate shape. (Fig. 4, A and B). We thus measured MT stability parameters within the spindle. First, we measured microtubule abundance within the inner-spindle. Despite the change in spindle shape, and in contrast to GTSE1-depleted cells, we could not detect a significant change in total inner-spindle tubulin fluorescence nor the total spindle volume occupied by microtubules in GTSE1^5xLID^ cells(Fig. 4 C; Fig. S5 A). We next specifically assayed kinetochore-MT attachment stability by treating cells with cold to depolymerize non-kMTs and quantifying the remaining MT intensity (Fig. 4, D and E). While depletion of GTSE1 led to 50% decrease in the tubulin fluorescence of cold resistant MTs, this defect was surprisingly not only rescued in GTSE1^WT^ cells, but also in the GTSE1^5xLID^ cells, indicating that GTSE1^5xLID^-GFP is able to stabilize kMTs. Finally, we quantified the stability of all growing MTs by inducing monopolar spindles with the Eg5 kinesin inhibitor STLC, and determining microtubule lengths by measuring distances between EB1-stained MT plus-ends and the centrosome. This assay revealed a clear reduction in both the length and abundance of microtubules in both GTSE1-depleted and GTSE1^5xLID^ cells (Fig. 4, F and G; Fig. S5 B). Thus, the CHC-GTSE1 interaction is dispensable for the maintenance of metaphase kinetochore-MT attachment stability, but not microtubule stability in general nor spindle morphology.

**Figure 4:**
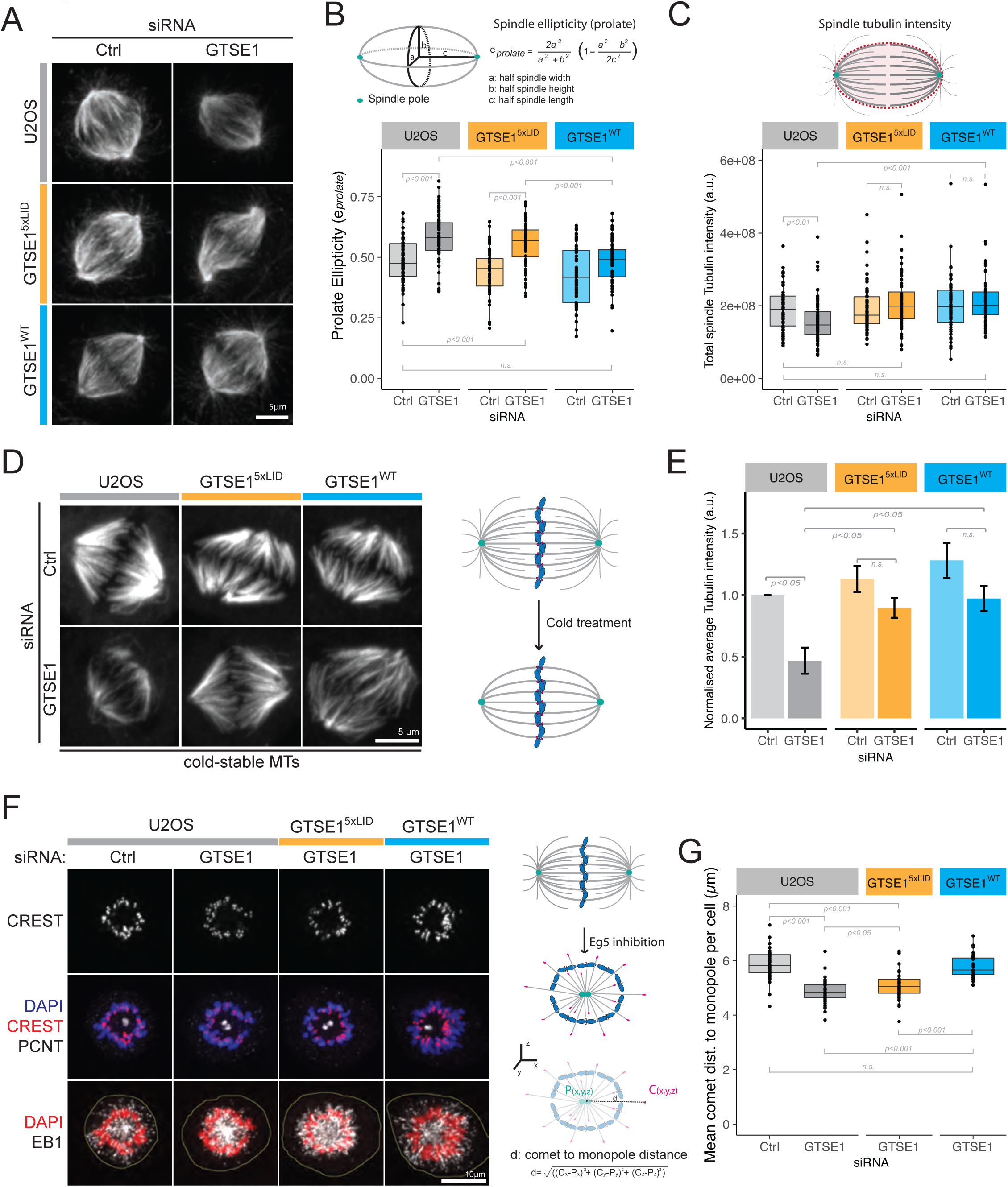
The CHC-GTSE1 interaction is required for non-kinetochore microtubule stabilization. **(A)** Immunofluorescence images of the mitotic spindle (α-tubulin staining) of U2OS, GTSE1^WT^ and GTSE1^5xLID^ cells, transfected with Control (Ctrl) or GTSE1 siRNA. The spindle prolate ellipticity **(B)** and the inner-spindle total tubulin fluorescence intensity **(C)** were measured in cells from (A) using 3D objects representing the spindles without astral microtubules (n>60 cells per condition, N=2 experiments pooled for representation and analysis, *p-values* from ANOVA followed by Tukey in (B) and Wilcoxon test in (C)). **(D)** Immunofluorescence images showing cold-resistant microtubules in U2OS, GTSE1^WT^ and GTSE1^5xLID^ cells, transfected with Ctrl or GTSE1 siRNA. G2-synchronized cells (RO3306) were released into mitosis for 55 minutes before cold treatment. α-tubulin staining is shown. **(E)** The total tubulin fluorescence intensity of cold-resistant microtubules was measured in cells from (D). The average total tubulin fluorescence intensity was calculated for each condition and normalized to U2OS Ctrl cells. Means of the normalized averages over 3 experiments are presented (+/-SE, n>151 cells per conditions, *p-values* from two-sided, unpaired t-test). **(F)** Immunofluorescence images showing STLC-induced monopolar spindles in U2OS, GTSE1^WT^ and GTSE1^5xLID^ cells transfected with Ctrl or GTSE1 siRNA. Cells were stained for EB1, PCNT, CREST and DNA. Dashed yellow lines indicate cell borders. **(G)** Mean growing microtubule length per cell measured as the EB1 comet to monopole distance following 3D reconstruction from experiment shown in (F) (n>36 cells per condition, N=3 experiments pooled for analysis and representation, *p-values* from Wilcoxon test). Numeric data in Table S1.

### Loss of CHC-GTSE1 interaction destabilizes astral microtubules

The above results suggest that the clathrin interaction with GTSE1 facilitates chromosome congression independently of kinetochore-MT attachment stability. How could stabilization of non-kMTs by the CHC-GTSE1 interaction facilitate chromosome congression? To address this question, we imaged both chromosome and spindle dynamics in live GTSE1^5xLID^ cells expressing mCherry ß-tubulin and with DNA labeled. Despite forming bipolar spindles in more than 95% of mitoses, these cells showed an increased mitotic duration and displayed misaligned chromosomes in 32.8% of mitoses (6.3% in control cells) (Fig. S6, A and B). Of the cells with misaligned chromosomes, 84.2% contained at least one chromosome transiently residing in the vicinity of one of the spindle poles before congression, reminiscent of chromosome behavior when congression is impaired, for example by loss of either dynein-mediated chromosome capture by astral MTs, CENP-E motor activity, or chromokinesin-mediated polar ejection forces (PEFs) (Schaar et al., 1997; Wandke et al., 2012; Li et al., 2007a).

One study has suggested that changes in MT dynamics following GTSE1 depletion cause a decrease in Aurora B activity/levels, which in turn leads to weaker PEFs, although the PEF defect does not appear to be the cause of the associated chromosome congression defect(Tipton et al., 2017). To ask whether the CHC-GTSE1 interaction controls PEFs or CENP-E function, we calculated the distances between kinetochores and poles of STLC-induced monopolar spindles, as has been previously described (Fig. 4 F) (Tipton et al., 2017; Barisic et al., 2014; 2015). We did not, however, observe any significant differences in the average kinetochore to monopole distances between U2OS control cells, GTSE1-depleted cells, and GTSE1^WT^ or GTSE1^5xLID^ cells depleted for GTSE1 (Fig. 4 F; Fig. S6 C). We thus analyzed the impact of GTSE1 depletion on Aurora B in our cells. Depletion of GTSE1 did not lead to any detectable decrease in Aurora B levels in mitotic cells (Fig. S6, D and E), nor in its activity as measured by quantifying histone H3 pSer10 on chromosomes (Fig. S6, F and G). Together these data indicate that defects in Aurora B level/activity, PEFs, or CENP-E function are unlikely to be the cause of the chromosome misalignment observed in U2OS cells following GTSE1 perturbation.

Because astral MTs can promote chromosome congression by capturing chromosomes located outside of the spindle area and facilitating their dynein-mediated poleward transport(Li et al., 2007b; Maiato et al., 2017), we specifically quantified the stability of astral MTs in GTSE1^5xLID^ cells. Depletion of GTSE1 decreased both the length and abundance of astral MTs(Fig. 5, A-D). While both astral MT parameters were restored in GTSE1^WT^ cells, they were clearly defective when the CHC-GTSE1 interaction was impaired in GTSE1^5xLID^ cells. This indicated a role of clathrin in stabilizing astral MTs. To confirm this, we measured astral MT lengths and abundance in CHC-depleted cells. Indeed, CHC depletion alone led to significantly fewer and shorter astral MTs (Fig. 5, E-G).

**Figure 5:**
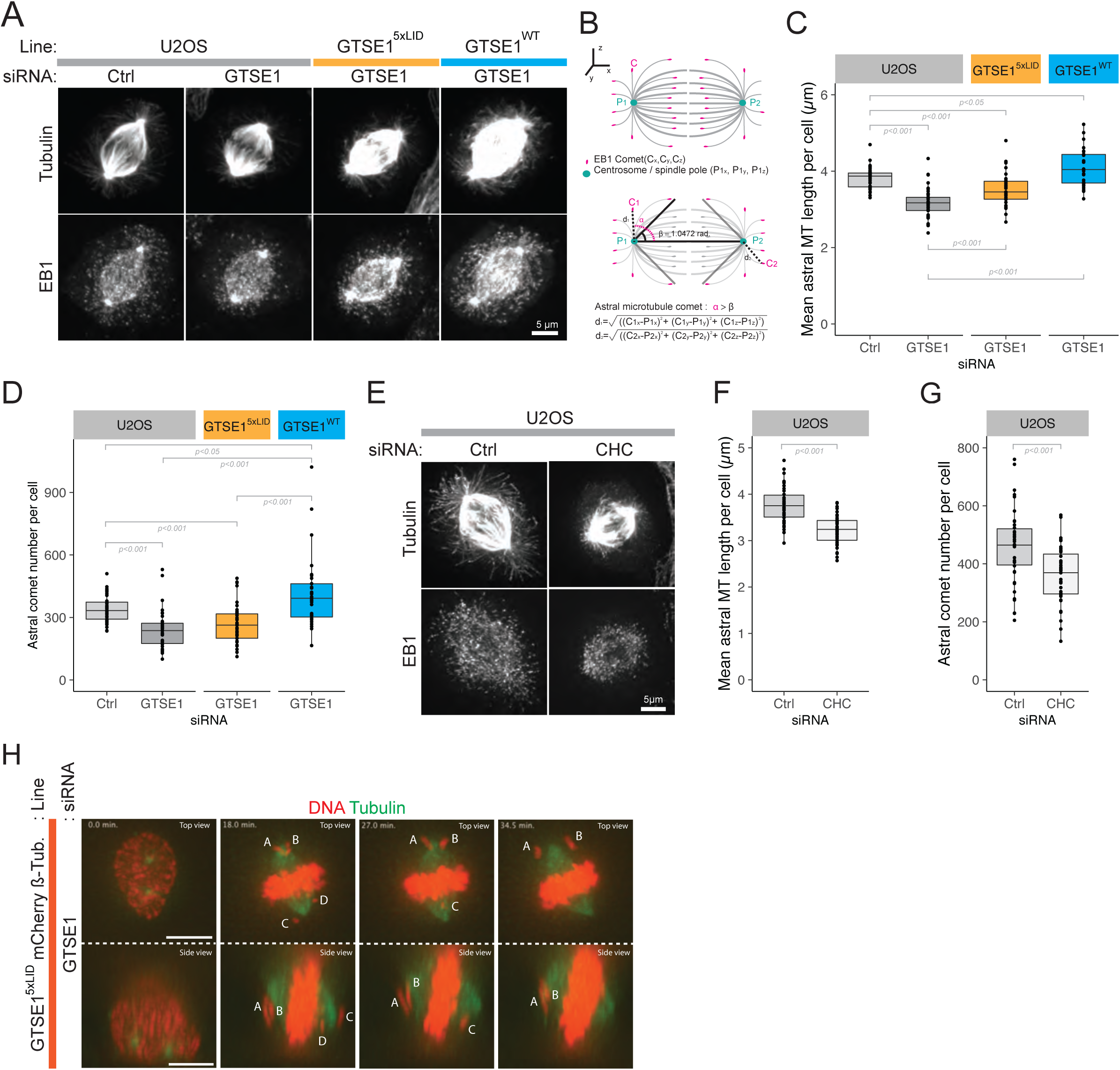
Astral microtubules are destabilized upon disruption of the CHC-GTSE1 interaction. **(A)** Immunofluorescence images illustrating astral length and abundance in U2OS, GTSE1^WT^ and GTSE1^5xLID^ cells, transfected with Control (Ctrl) or GTSE1 siRNA. EB1 and α-tubulin staining are shown. **(B)** Scheme illustrating the quantification method used to measure astral MT length and abundance in 3D reconstruction of cells from (A). Astral microtubule length **(C)** was measured as the distance between astral EB1 comets and the closest spindle pole. The mean astral microtubule length per cell is presented. The astral microtubule abundance **(D)** was measured as the number of astral EB1 comets per cell. (N=3, n>30 cells per condition pooled for analysis and representation, *p-values* from Wilcoxon test). **(E)** Immunofluorescence images illustrating astral microtubule length in U2OS cells transfected with Ctrl or CHC siRNA (114 h). α-tubulin and EB1 staining are shown. Quantification of astral microtubule length **(F)** and abundance **(G)** in cells from (E) as in (C) and (D), respectively (n>41 cells per condition, N=4 exp. pooled for representation and analysis, *p-value* from ANOVA and Wilcoxon test, respectively) **(H)** Stills from live cell imaging of GTSE1^5xLID^ cells expressing a mCherry ß-tubulin BAC transgene and with DNA stain, transfected with GTSE1 siRNA. Top and side view from a 3D reconstruction are presented. Scale bar: 10 µm. Letters indicate misaligned chromosomes. Notice misaligned chromosomes remaining outside of inner spindle mass when viewed in 3D. Numeric data in Table S1.

We then asked whether the behavior and attachment status of misaligned chromosomes occurring upon disruption of the CHC-GTSE1 interaction were consistent with a defect an astral MT capture. Closer inspection of misaligned chromosome positions and trajectories in three dimensions in mCherry-ß-tubulin GTSE1^5xLID^ cells depleted for endogenous GTSE1 revealed that most misaligned chromosomes resided outside of the spindle (i.e. where astral MTs should normally be present) for an extended period of time, before being able to congress to the metaphase plate (Fig. 5 H, Supplementary Video 1). Closer analysis of misaligned chromosomes in GTSE1^5xLID^ cells (39 chromosomes from 16 cells) revealed that most (73%, 14/19) kinetochore pairs located clearly outside of the spindle showed Mad1 staining on both kinetochores, indicating they both lacked MT attachment (Chen et al., 1998)(Fig. S6 H upper lane). In contrast, this was observed for only 22% (2/9) of misaligned chromosome pairs located clearly within the spindle (Fig. S6 H lower lane), indicating that the majority of these had established a kinetochore-MT attachment and were likely congressing.

Together these data show that most misaligned chromosomes in GTSE1^5xLID^ cells are located outside of the spindle poles and lack robust KT-microtubule attachments. This is consistent with a scenario in which astral MTs are too short and sparse to efficiently allow chromosome capture and/or congression. Thus, the CHC-GTSE1 interaction predominantly facilitates chromosome congression through stabilization of non-kinetochore-MTs.

### Clathrin promotes chromosome congression and timely mitosis by antagonizing MCAK activity via GTSE1 recruitment

MCAK activity regulates astral MT stability, and we previously showed that GTSE1 binds and inhibits MCAK (Bendre et al., 2016; Srayko et al., 2005; Rizk et al., 2009). To test whether the phenotypes observed following disruption of the CHC-GTSE1 interaction resulted from excess MCAK activity, we treated GTSE1^5xLID^ cells with GTSE1, MCAK, or GTSE1 and MCAK siRNA and monitored the time between NEB and anaphase onset (Fig. 6 A, Fig. S7 A). Strikingly, codepletion of MCAK and endogenous GTSE1 in GTSE1^5xLID^ cells eliminated the increased time to anaphase onset associated with loss of the CHC-GTSE1 interaction. This was also observed in GTSE1^5xLID^ cells expressing mCherry ß-tubulin (Fig. S6 B). Furthermore, the percentage of mitotic cells displaying misaligned chromosomes was dramatically reduced, from 32.8% in GTSE1 depleted cells to 11.6% in GTSE1 and MCAK co-depleted cells (Fig. S6 B). We then asked if the MT stability defect observed upon disruption of the CHC-GTSE1 interaction also depends on MCAK activity. Measurement of MT length in GTSE1^5xLID^ cells treated with STLC showed that the decrease in MT length observed upon depletion of GTSE1 was alleviated upon co-depletion with MCAK (Fig. 6, B and C). These data are consistent with the MT stability and chromosome congression defects following disruption of the CHC-GTSE1 interaction arising from excess MCAK activity.

**Figure 6:**
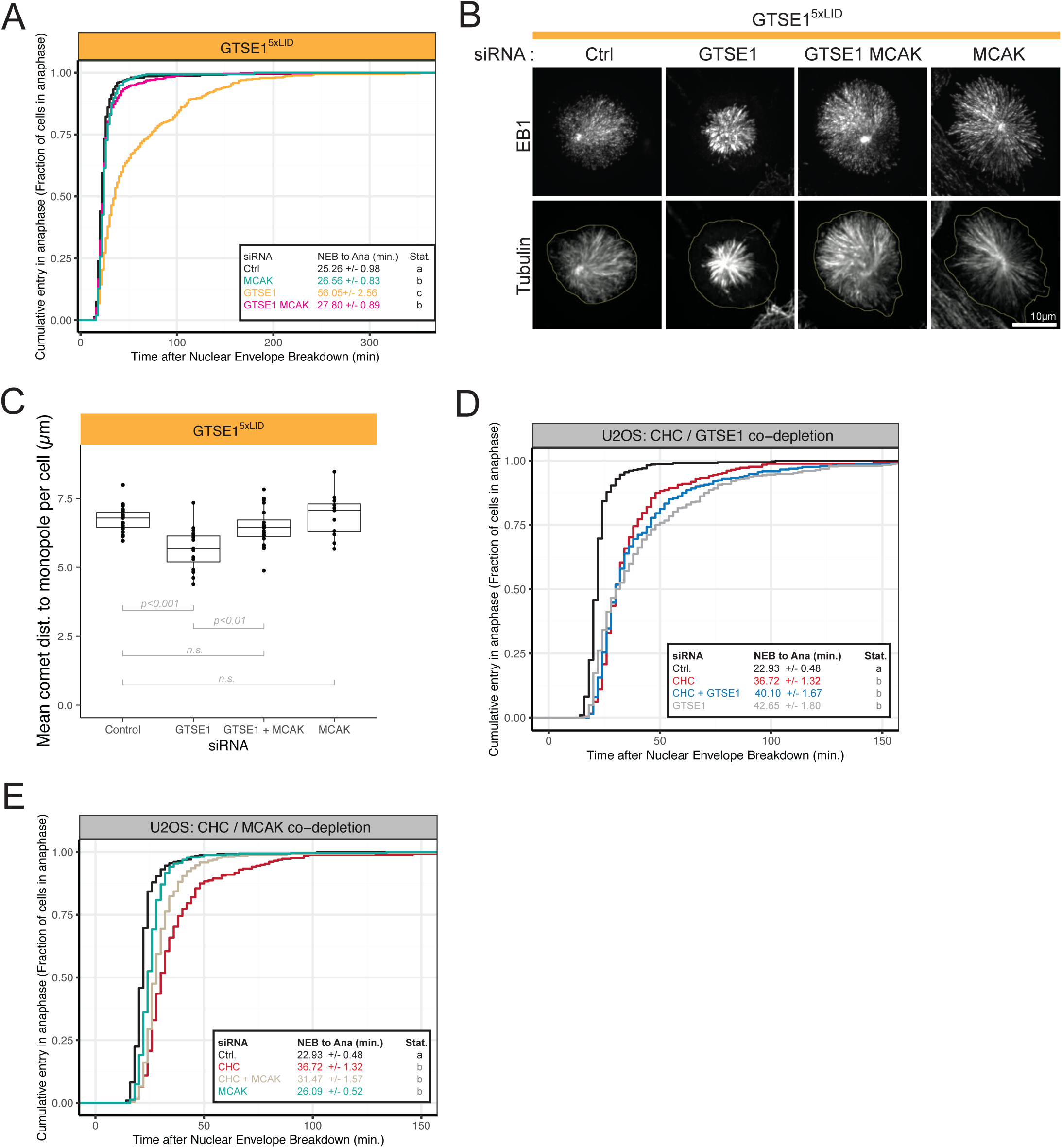
Clathrin promotes chromosome congression and timely mitosis by antagonizing MCAK activity via GTSE1 recruitment. **(A)** Depletion of MCAK restores anaphase onset delay defect in GTSE1^5xLID^ cells. Fraction of cells that have entered anaphase as a function of the time (min) after nuclear envelope breakdown (NEB) in GTSE1^5xLID^ cells transfected with control (Ctrl), GTSE1, MCAK, or GTSE1 + MCAK siRNA(s). The mean time between NEB and anaphase onset is indicated in the inset (+/-SE, n>311 per condition over N=4 experiments pooled for analysis and representation, *p-value* from Wilcoxon test). **(B)** Immunofluorescence images showing STLC-induced monopolar spindles in GTSE1^5xLID^ cells, transfected with Ctrl, GTSE1, MCAK, or GTSE1 + MCAK siRNA(s). Tubulin and EB1 staining are shown. Dashed yellow lines indicate cell borders. **(C)** Measurement of growing microtubule length (EB1 comet to monopole distance) in 3D reconstruction of cells from (B). The mean microtubule length per cell is presented. (Ctrl n=19, GTSE1 n=20, GTSE1 + MCAK n=20, MCAK n=12, N=1, *p-value* from ANOVA followed by Tukey). **(D)** CHC and GTSE1 depletion in U2OS cells lead to similar and non-additive mitotic delays. Data plotted as described in (A). **(E)** Co-depletion of MCAK partially rescues the mitotic delay induced by CHC depletion. Data plotted as in (A). Data in (E) and (F) were obtained concomitantly (Ctrl and CHC siRNAs conditions are shared, n>254 over N= 3 experiments pooled for analysis and representation). U2OS cells were treated with control (Ctrl), CHC, GTSE1, MCAK, CHC and GTSE1 or CHC and MCAK siRNA(s). Plots in (E), and (F) are cropped on the time abscissa at t=150 min for representation purposes. In the “Stat.” columns, conditions with the same letter are not statistically different. (Wilcoxon test, *p<0.05*). Scale bar: 10 µm. Numeric data in Table S1.

Finally, we studied the contribution of GTSE1 to clathrin’s role in spindle assembly and chromosome alignment, using the time between NEB and anaphase onset as a measure of the pathway functionality. Depletion of GTSE1 or CHC resulted in very similar defects in mitotic timing, consistent with their similar effects on astral microtubule stability. Strikingly, combining loss of GTSE1 and CHC did not lead to a mitotic delay greater than depletion of either protein alone (Fig. 6 D; S7 B). These observations are consistent with GTSE1 acting downstream of CHC and significantly contributing to clathrin’s function in this process. We thus asked if the delayed mitosis after loss of CHC was also dependent on MCAK activity. Indeed, U2OS cells co-depleted for CHC and MCAK took significantly less time to transit from NEB to anaphase onset than U2OS cells depleted for CHC alone (Fig. 6 E, Fig. S7 B). Altogether, these observations indicate that a substantial part of CHC function in promoting MT stability, chromosome congression, and timely mitosis is to directly recruit the clathrin effector GTSE1 to antagonize MCAK activity.

## DISCUSSION

Clathrin’s importance during mitosis for chromosome alignment has only recently been appreciated. The role of spindle-associated clathrin for this function has generally been assumed, based on studies to date, to be to specifically stabilize and organize k-fibers. Here we have shown that clathrin functions to stabilize non-kMTs, such as astral MTs, and that this is an important mechanism by which clathrin promotes chromosome congression. This function of clathrin is mediated via clathrin adaptor-like interactions of GTSE1 with the CHC NTD, demonstrating for the first time a role for this well-characterized endocytic interaction scheme in mitosis. In contrast to clathrin adaptor proteins that utilize this interaction to recruit clathrin to sites of coat formation, clathrin on the spindle utilizes it to recruit GTSE1 and thus stabilize microtubules via GTSE1’s ability to inhibit MCAK. Thus, a major mechanism by which clathrin stabilizes MTs is ultimately inhibition of MCAK.

A schematic model of the clathrin/TACC3 complex based on studies to date is illustrated in Fig. 7. In contrast to TACC3 and CHC(Hubner et al., 2010; Lin et al., 2010; Fu et al., 2010; Booth et al., 2011), GTSE1 does not appear to be required for the localization of CHC or TACC3, yet is dependent on complex integrity for its localization to the spindle. GTSE1 thus likely functions as an “effector” of this complex. Therefore, many of the phenotypes attributed to complex disruption (i.e depletion of CHC, TACC3, or PI3K-C2*α*) may partly result from delocalization of GTSE1 from spindles and loss of MCAK inhibition.

**Figure 7:**
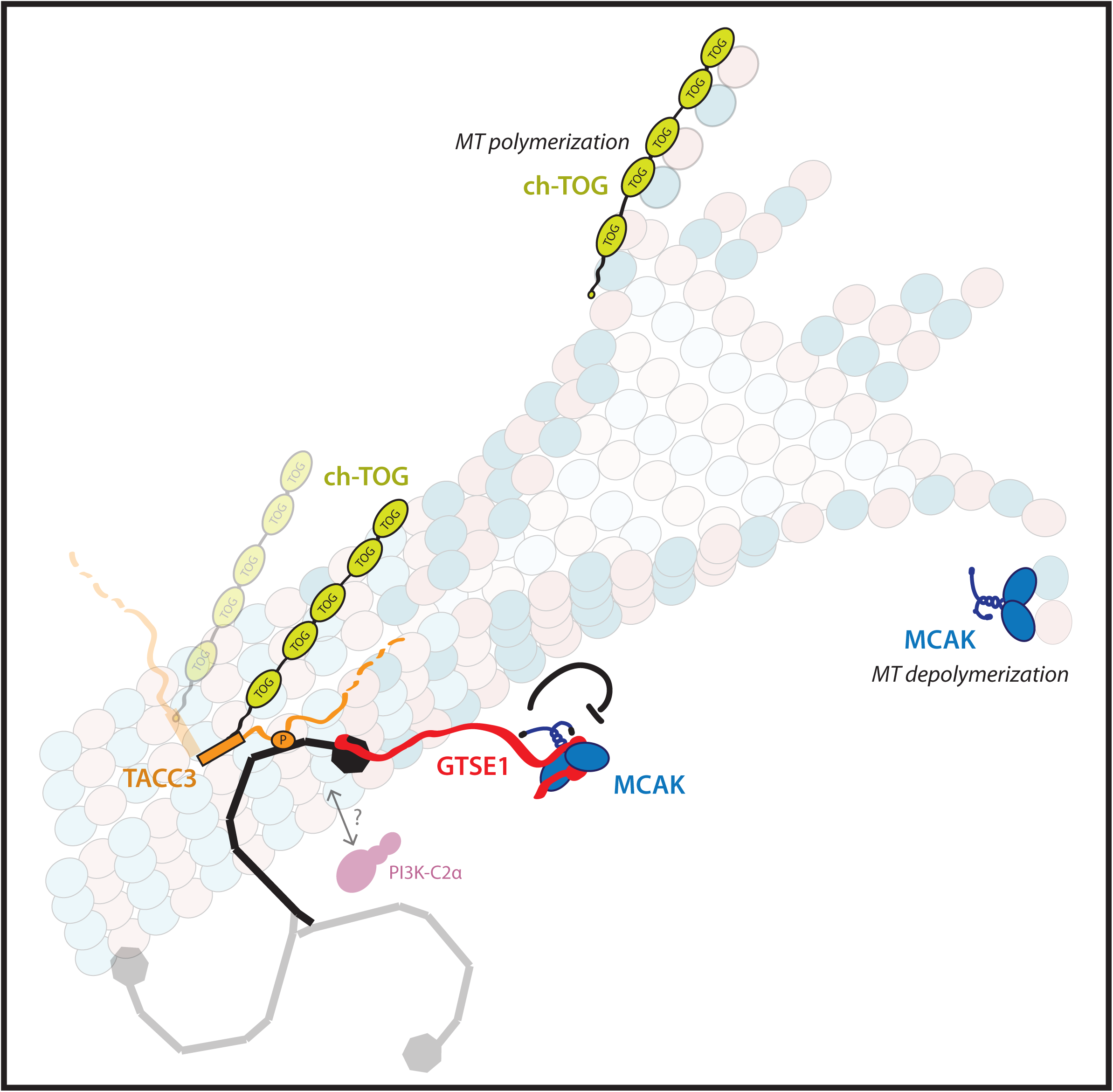
Model of the clathrin/TACC3 complex on microtubules. Schematic model of the clathrin/TACC3 complex on a microtubule based on studies to date. GTSE1, CHC, PI3K-C2a, TACC3, ch-TOG, MCAK and a microtubule are represented. The Aurora A dependent phosphorylation of S558 on TACC3 is represented as a P. TACC3 is also bound to the microtubule polymerase ch-TOG. The CHC N-terminal domain recruits GTSE1 to inhibit the microtubule depolymerase MCAK. It is unknown whether PI3K-C2a and GTSE1 can interact with a single CHC on the spindle. Similarly, as no information on the stoichiometry of the members of the complex are available, potential multimers are shaded. Relative protein sizes are roughly conserved.

Interestingly, while CHC and TACC3 homologs are conserved among eukaryotes(Peset and Vernos, 2008; Gergely, 2002), published reports of CHC and TACC3 interacting on the mitotic spindle have been limited to vertebrate model systems (human, mouse, chicken, and Xenopus), and it has been proposed that this function of clathrin may have arisen later in evolution(Gulluni et al., 2017; Royle, 2013; Hubner et al., 2010; Lin et al., 2010; Fu et al., 2010; Burgess et al., 2015). Our finding that homologs of human GTSE1 containing the important C-terminal domain only exist in vertebrates raises the question of whether the appearance of GTSE1 (and its interaction with CHC) may be evolutionarily linked to clathrin localization to the spindle and its MT stabilizing function during mitosis. However, we find that TACC3 residues identified as required for interaction with CHC (Hood et al., 2013; Burgess et al., 2015; 2018) are more distantly conserved, suggesting that a CHC-TACC3 complex on the spindle may have a more ancient origin; Fig. S8). Nevertheless, given that the C-terminal domain of GTSE1 is restricted to vertebrates, the GTSE1-dependent MT stabilization role of the CHC/TACC3 complex likely has a more recent evolutionary origin.

With our finding that CHC recruits GTSE1, the clathrin/TACC3 complex becomes a hub incorporating regulation of two major effectors of MT dynamics, ch-Tog/XMAP215 and MCAK, to stabilize microtubules(Gard and Kirschner, 1987; Vasquez et al., 1994; Walczak et al., 1996; Desai et al., 1999; Tournebize et al., 2000; Kinoshita et al., 2001). We have shown that GTSE1 inhibition of MCAK provides the major contribution of this complex’s ability to stabilize astral MTs and promote chromosome alignment, as co-depletion of GTSE1 and CHC together does not worsen chromosome alignment defects more than either single depletion, and depleting MCAK concurrently with CHC alleviates the severity of the CHC depletion defect (Fig6 E, F). Thus, while it has been assumed that the ability of TACC3 to antagonize MCAK activity at spindle poles to regulate astral microtubule stability is via promoting ch-Tog polymerization activity(Kinoshita et al., 2005), these results indicate an additional more direct mechanism: TACC3 in complex with CHC directly recruits the MCAK-inhibitor GTSE1. Determining the precise contribution of CHC/TACC3 complex-associated ch-TOG to microtubule stability has proven difficult to assess, as ch-TOG has multiple roles independent of this complex in spindle assembly, most ch-TOG protein resides at centrosomes and kinetochores independently of CHC/TACC3 complexes, and a TACC3/ch-TOG subcomplex resides on MT plus-ends independent of CHC(Al-Bassam and Chang, 2011; Booth et al., 2011; Gergely et al., 2003; Gutierrez-Caballero et al., 2015; Lin et al., 2010).

Our results are consistent with multiple mechanistic roles for clathrin operating concurrently during mitosis. Evolutionary analysis of CHC, TACC3, and GTSE1 suggests that a mitotic CHC/TACC3 complex may have existed before vertebrates, while GTSE1 interaction with CHC likely originated in vertebrates, consistent with separate roles of the CHC/TACC3 complex. In addition, while co-depletion with MCAK restores mitotic duration in cells where the CHC-GTSE1 interaction is prevented, in CHC-depleted cells this effect is partial, implicating another potential role in mitosis. One potentially GTSE1-independent role of the CHC/TACC3 complex is the organization of k-fiber MTs via the inter-MT “mesh” observed in electron micrographs(Nixon et al., 2015). This role has been used to explain observations that loss of clathrin leads to chromosome alignment defects and less MTs within k-fibers, and that excess TACC3 leads to more MTs in k-fiber cross-sections, many of which are less parallelly oriented, and more “mesh” interconnections(Booth et al., 2011; Cheeseman et al., 2013; Nixon et al., 2015). Physical bridges/mesh between MTs along the length of k-fibers may explain “stabilizing” MTs within a k-fiber bundle from an organizational perspective. However, it is not clear how they would affect the dynamics/stabilization of kinetochore MTs within k-fibers, whose plus-ends are by definition at the kinetochore. In fact, no kinetochore-MT attachment defect was observed upon removal of TACC3/CHC from metaphase spindles or overexpression of TACC3, although k-fiber MT number/organization changed(Cheeseman et al., 2013; Nixon et al., 2015). Perhaps the changes in MT numbers within k-fibers observed after CHC or TACC3 perturbation by EM represent loss of non-kMTs present within the k-fibers, not detectable from fluorescence microscopy assays. The CHC-GTSE1-MCAK pathway may function to stabilize such a MT population, such as augmin-dependent microtubules(David et al., 2019), and/or stabilize damaged MTs within k-fibers against catastrophe(Aumeier et al., 2016). Alternatively, or additionally, this could be precisely a role of spindle-localized ch-TOG. Either way, our data support the earlier conjecture that the CHC/TACC3 complex has dual roles on the spindle to organize MTs, as well as to inhibit MT catastrophes (i.e. stabilize MTs), the latter albeit not only within, but also outside of k-fibers(Booth et al., 2011). It will be interesting to see whether loss of GTSE1 affects k-fiber composition of MTs or the “mesh”, as well as the effect of specifically perturbing spindle associated, CHC/TACC3-recruited ch-TOG.

Our structures of the CHC-GTSE1 interface allow us to compare this interaction to that of CHC-adaptor interactions that function in endocytosis. GTSE1 is able to make multiple contacts with the CHC NTD. Site 3 on the CHC NTD clearly binds to motif D on GTSE1. Interestingly, the GTSE1 residues involved in this interaction with Site 3 fit the consensus of a CBM and not the reported Site 3-binding motif [LI][LI]GXL(Kang et al., 2009; Haar et al., 2000), although a sequence similar to this in GTSE1 (VVGQL) adjacent to motif D makes contacts nearby. The interacting residues of motif D (LIDL) and their position within Site 3 are in fact very similar to what has been recently observed with short peptides(Muenzner et al., 2016). Our data thus show that CBMs can indeed interact with more than a single site on the CHC NTD, even in a larger protein domain context, supporting reports using small peptides(Zhuo et al., 2015; Muenzner et al., 2016). Additionally, our work suggests that different CBMs within a polypeptide stretch, commonly observed in adaptor proteins, could interact with a single CHC NTD, and enhance binding affinity. The spacing of the GTSE1 motifs would allow a bidentate binding to the CHC NTD with motif D in site 3 and motif B in site 1. While the expanded repertoire of clathrin adaptor proteins and motifs that function in endocytosis results in functional redundancy of CHC adaptor-binding sites for transferrin endocytosis, this was not observed to be the case for the CHC mitotic role, where disruption of Site 1 alone, but not Site 4, perturbs function(Hood et al., 2013). Thus there is likely a more limited and/or specific set of interactions with the CHC adaptor-binding sites required on the spindle. Whether our observed GTSE1 binding preferences explain this difference alone, or whether interactions of additional proteins with CHC adaptor sites are also important for mitosis, will be important to establish. Our elucidation of the GTSE1-CHC interaction interface raises the question of whether GTSE1 also functions as a clathrin adaptor in vesicle trafficking. Interestingly, CHC and GTSE1 also interact in interphase (unpublished data), and tethering the C-terminus of GTSE1 to membranes can recruit clathrin and initiate endocytosis, suggesting it may(Wood et al., 2017).

Our results also indicate a separation of function for GTSE1 roles in microtubule stabilization. While depletion of GTSE1 from U2OS cells leads to defects in the stability of astral MTs, inner-spindle MT and kinetochore-MT attachment, GTSE1^5xLID^ that cannot interact with clathrin only exhibits defects in the former, even though apparently completely delocalized from the spindle(Bendre et al., 2016). GTSE1 therefore appears to have the ability, independent of clathrin, to stabilize kinetochore-MT attachment. This could hypothetically function via inhibition of centromere-associated MCAK. However, we have been unable to observe specific localization of GTSE1 to centromeres. Because GTSE1 levels in U2OS and other cancer cell lines are extremely elevated as compared to untransformed cells(Scolz et al., 2012), the observed stabilization of kinetochore-MT attachments by GTSE1 may only occur when GTSE1 is overexpressed, and not represent a normal function of the protein. Consistent with this idea, U2OS cells have approximately a 2-fold increase in kMT stabilization as compared to “normal” RPE-1 cells, while removal of GTSE1 from U2OS cells in our hands results in approximately 2-fold decrease in kMT stabilization(Bakhoum et al., 2009; Bendre et al., 2016).

The mechanisms by which clathrin operates in mitosis have been somewhat of an enigma. Here we provide a direct mechanistic link to microtubule stabilization, whereby adaptor interaction sites in the CHC NTD directly recruit a “clathrin effector”, the microtubule stabilizing protein GTSE1, to inhibit MCAK activity and stabilize centrosomal microtubules. Studies on GTSE1 suggest that its accurate expression and regulation is not only important for chromosome alignment and spindle positioning, but also for avoiding chromosomal instability and potentially tumour progression(Bendre et al., 2016; Tipton et al., 2017; Scolz et al., 2012).

## ACKNOWLEDGEMENTS

We thank A. Musacchio for purified Aurora B and Cdk1 kinases and anti-MAD1 antibody, H.M. Shih for CHC expression constructs, S.J. Royle for a full-length CHC plasmid, K. Kinoshita for phospho-TACC3 antibody, L. Wordemann for anti-MCAK antibody, and A.A. Hyman for mCherry-ß-tubulin and CHC-GFP BACs. We thank G. Vader, R. Cardoso da Silva, M. Pesenti, S. Maffini, and J. Almario for comments on the manuscript. We thank S. Maffini for help with microscopy. We thank the beamline staff of X10SA at the Swiss Light Source Paul Scherrer Institute, Villigen, CH, for support, and our colleagues of MPI Dortmund and the TU Dortmund for help with the data collection.

This work was supported by the Max Planck Institute of Molecular Physiology, a Worldwide Cancer Research project grant 16-0093 to A.W. Bird, and the Deutsche Forschungsgemeinschaft (DFG, German Research Foundation) – Projektnummer 394694869

## AUTHOR CONTRIBUTIONS

A.R., Y.L., D.S., A.P., H.C.T, M.H., I.V, and A.B. conceived and designed experiments.

A.R. performed most cellular assays, image quantification, analysis, and statistics.

Y. L. and P.B. constructed and purified protein fragments and mutants, performed all pull-down assays and Y2H assays, and purified TACC3, and CHC protein constructs

A.P. and I.V. performed crystallization experiments and analysis

D.S. performed live cell imaging and analyses of cellular Aurora B expression and activity

P.W. purified Aurora A kinase and GTSE1-GFP proteins

A.H. quantified CHC localization

H.C.T. performed AUC analysis

M.H. performed sequence/evolutionary analyses

N.S. generated cell lines

S.B. purified GTSE1 protein

F.M. and T.B. performed mass spectrometry experiments and analysis

A.R. and A.B. wrote the manuscript.

## METHODS

### Cloning and plasmids

The full-length clathrin cDNA was provided by SJ Royle (CMCB, Warwick medical school). The full-length GTSE1, clathrin and TACC3 cDNAs were subcloned into pFastbac vectors with a N-terminal GST tag and a Precision protease cleavage site between the GST and the protein of interest. pGEX4T1-CHC-1-330, 331-1163 and 996-1654 were gifts from HM Shih (Academia Sinica, Taiwan). Clathrin fragments were amplified by PCR and subcloned into pGEX-6P1 vector and CHC-1-642 was subcloned into pET28a with His tags. Coding sequences for human GTSE1 and its variants were amplified using PCR and cloned into a plasmid vector. GTSE1-C 639-739; GTSE1-C1 (652-687); GTSE1-C2(672-708); GTSE1-C3 (693-739) were cloned into pGEX-6P1. GTSE1 1-460; GTSE1 463-739; GTSE1 639-739 were cloned into pET28a in frame with a His Tag. Mutations on clathrin or GTSE1 were generated by site directed mutagenesis. Clathrin mutants for the adaptor binding sites on the NTD were obtained by introducing the following mutations: T87A, Q89A, K96E and K98E (S1M), Q152L and I154Q (S2M), and R188A and Q192A (S3M). The CBM on GTSE1 were mutated by introducing the following alanine substitution: LID_(664-666)_ to AAA (motif A), LID_(669-671)_ to AAA (motif B), LID_(689-691)_ to AAA (motif C), LID_(713-715)_ to AAA (motif D), IDL_(714-716)_ to AAA (motif D), LIDL_(713-716)_ AADA (motif D), and E mutant LIQ_(720-722)_ to AAA (motif E). CHC 1-364 and GTSE1 (661-726, C4 or 653-719, C5) were subcloned into pGEX-2rbs, a dicistronic derivative of pGEX6P generated in house, which allowed two genes to be translated from the same transcript. In this construct clathrin has a N-terminal GST tag while GTSE1 has no tag.

The GTSE1-GFP BAC resistant to the siRNA targeting the sequence GATTCATACAGGAGUCAAA is described in Scolz *et al*., 2012 (Scolz et al., 2012). To obtain the GTSE1^5xLID^-GFP BAC, the point mutations LID(664-666)AAA, LID(669-671)AAA, LID(689-691)AAA, LID(713-715)AAA, and LIQ(720-722)AAA were introduced into the GTSE1-GFP BAC through two successive rounds of counterselection recombineering based on Redβ and Redγ expression(Bird et al., 2011). The mCherry ß-tubulin BAC (MCB6696, HS.E044.J01 backbone) and CHC-GFP BAC (MCB4422, RP11-661B17 backbone) were kind gifts from the Hyman Lab.

## Antibodies

Antibodies used: goat anti-GFP (MPI Dresden; described in Poser *et al*., 2008), rabbit anti-GTSE1 (custom generated; described in Scolz *et al*., 2012), rabbit anti-TACC3 (sc-22773, Santa Cruz Biotechnology), mouse anti-α-tubulin (DM1α, Sigma-Aldrich), rat anti-tubulin (YL1/2, Santa Cruz Biotechnology), mouse anti-CHC (X22, ab2731, Abcam), rabbit anti-GFP (ab6556, Abcam), rat anti-EB1 (KT-51, Absea Biotechnology), mouse anti-MCAK/Kif2C (1G2, Abnova Corp.), human nuclear antibodies to nuclear antigens centromere autoantibody (CREST) (CS1058, Europa Bioproducts Ltd), rabbit anti-pericentrin (PCNT) (ab4448, Abcam), mouse anti-c-myc (Oncogene EMD Millipore), rabbit anti-Aurora kinase B (ab2254, Abcam), mouse anti-phospho-histone-Ser10 (H3pS10) (ab14955, Abcam), mouse anti-His (Penta-His, 34660, Qiagen), rabbit against phospho-TACC3 (gift of K. Kinoshita), mouse anti-MAD1 conjugated to Alexa488 (gift of A. Musacchio, MPI-Dortmund), donkey anti-human antibodies conjugated to Cy5 or Texas red (Jackson ImmunoResearch laboratories; #709-175-149 and #709-075-149), donkey anti-rat Alexa488 (Bethyl; A110-337D2, donkey anti-rat Alexa594 (Bethyl; A110-337D4), donkey anti-rabbit Alexa488 (Therofisher; A-21206), donkey anti-rabbit Alexa594 (Bethyl; A120-208D4), donkey anti-rabbit Alexa650 (Bethyl; A120-208D5), donkey anti-mouse Alexa488 (Bethyl; A90-337D2), donkey anti-mouse Alexa594 (Bethyl; A90-337D4), donkey anti-mouse Alexa647 (Invitrogen; A31571), donkey anti-goat Alexa488 (Jackson Immunoresearch; 705 545 147), donkey anti-goat HRP (Santa Cruz; SC-2020), sheep anti-mouse HRP (Amersham; NXA931-1ml), donkey anti-rabbit HRP (Amersham; NXA934-1ml).

## Protein expression and purification

The full-length GTSE1, clathrin and TACC3 were produced from Tnao38 insect cells as described previously(Bendre et al., 2016). In brief, cell pellets were harvested at 72 to 96 hours after infection and lysed into a buffer containing 50 mM HEPES, pH 8.0, 300 mM NaCl, 5% glycerol and protease inhibitors. The supernatant was cleared by centrifugation and applied onto a glutathione resin. To remove the GST tags, purified proteins were incubated with GST-Precision protease overnight at 4 °C, and applied onto a Superose 6 10/300 column. To obtain the clathrin-GTSE1 complex for crystallization, CHC 1-364 and GTSE1 (661-726, C4 or 653-719, C5) were co-expressed in E. *coli* BL21 (DE3) cells. Cells were lysed in GST-binding buffer (25 mM HEPES pH7.5, 300 mM NaCl, 1 mM EDTA, 5% glycerol, 0.5% Trion X-100 and protease inhibitors [Serva]). The bacterial lysate was first applied on GST affinity column, and the GST bound fraction was eluted in buffer A pH8.0 (50 mM HEPES, 300 mM NaCl, 5% glycerol and 20 mM reduced glutathione) and subjected to size-exclusion chromatography using a Superdex 75 16/600 column. To cleave the GST tag, fractions containing the complex were incubated with GST-PreScission protease overnight at 4 °C. GST and GST-PreScission protease were removed by applying the fractions onto a Superdex 75 16/600 column attached to a GST column and equilibrated in a buffer containing 20 mM Tris pH 7.0 and 50 mM NaCl. The clathrin-GTSE1 complexes were concentrated to approximately 30 mg/ml for the crystallization trail *(See Supplementary informations clathrin-GTSE1 Xray methods)*. Truncated fragments of clathrin or GTSE1 with GST or His tag were expressed in E. *coli* BL21 (DE3) cells and further purified onto glutathione, Ni-NTA or amylose resin. GTSE1-GFP was purified as described in *Scolz et al.,* 2012(Scolz et al., 2012). pET28 containing N-terminally 6xhis-tagged Aurora kinase A was transformed into Rosetta (DE3) cells with Kanamycin and Chloramphenicol selection. One liter of culture was grown to OD600 1.0 at 37 °C and cooled to 18 °C. 200 µL of 1M IPTG was added and the culture was grown overnight (18 hours) at 18 °C. Cells were pelleted and resuspended in 30 mL of lysis buffer (50 mM Tris-HCl pH7.2, 500 mM NaCL, 0.1% TritonX-100, 5% glycerol) and frozen in liquid nitrogen. Pellet was then thawed on ice with immediate addition of protease inhibitors (1 complete tablet [Roche], 40 µL peptstatin at 10mg/ml, 40 µL PMSF at 10 mg/mL) followed by lysozyme to 0.5 mg/mL final.

Suspension was sonicated for 30 seconds at 50% amplitude on ice. The lysate was cleared by centrifugation at 40,000 rpm for 15 min using an MLA-80 rotor. Imidazole was added to the collected supernatant to 10 mM final. The cleared lysate was loaded onto a 5 mL HisTrap column (GE life sciences) and washed with buffer (25 mM Tris-HCl pH 8.0, 300 mM NaCl, 20% glycerol) containing 10 mM Imidazole (50 mL) and 30 mM Imidazole (25 mL). Protein was eluted with the same buffer containing 300 mM Imidazole. Peak fractions were pooled and run on a Superdex 200 16/60 column with size exclusion buffer (50 mM Tris pH 7.5, 300 mM KCl). Peak fractions were pooled, adjusted to 1 mM DTT, 10% glycerol, aliquoted and snap frozen in liquid nitrogen.

## In vitro protein phosphorylation

Proteins of interest were phosphorylated by Aurora A, Aurora B or Cdk1 using a molar ratio 1:100 (kinase: protein) overnight on ice in buffer supplemented with 10 mM MgCl_2_, 1 mM sodium orthovanadate and 2 mM ATP. The phosphorylation reaction was stopped by adding 5 μM RO-3306 (Cdk1 inhibitor) or 500 nM MLN8054 (Aurora A inhibitor) for 10 min on ice. Protein phosphorylation was confirmed using Pro-Q Diamond staining (Invitogen). Purified Aurora B and Cdk1 were gifts from A. Musacchio.

## GST pull-down assays

In Brief, 1-5µg GST-tagged proteins were immobilized on GST resins and incubated with 2-5 μM proteins of interest were added in GST-binding buffer (25 mM HEPES pH7.5, 300 mM NaCl, 1 mM EDTA, 5% glycerol and 0.5% Trion X-100) for 1∼3 hour at 4 °C. The reactions were washed with GST binding buffer, and analyzed by SDS-page or Western blotting.

## Yeast-two hybrid

Clathrin 1-330 was cloned into pGAD vector and GTSE1 463-739 WT or CBM mutants were in-frame inserted into pGBD vector. Plasmids were transformed into *S. cerevisiae* cells using the lithium acetate, single-strand carrier DNA, polyethylene method(Gietz and Woods, 2002). Cells containing both two-hybrid plasmids were selected on a double dropout plate (SD-ura-leu). Single colony was picked and grown in the double dropout medium till O.D 0.6. Cells were serially diluted and dropped on a double dropout (SD-ura-leu) and triple dropout (SD-ura-leu-his) plates. The interaction strength was examined on the triple dropout plate.

## Analytical Ultra Centrifugation

Bacterial expressed CHC (1-364) was purified as GST fusion protein (using standard protocol and standard buffer (Tris 25mM pH 7.5, NaCl 100mM, EDTA 2mM and DTE 1mM)) and GST tag was cleaved using GST Precision protease. Further tag was removed by performing reverse GST purification followed by gelfiltration using superdex 75 column using standard buffer. Bacterial expressed GTSE1(639-739)-6xHis protein was purified as N-terminal MBP fusion protein (using standard protocol and standard buffer (Tris 25mM pH 7.5, NaCl 50mM, β-ME 2mM)) and MBP-tag was cleaved using GST-Precision protease followed by Ni-IMAC chromatography using standard protocol (wash and elution buffer containing 30mM and 600mM Imidazole respectively in standard buffer). Later buffer was exchanged on centricon concentrator against standard buffer. For molecular size determination, two proteins were mixed in equal molar ratio in 200μl volume and injected on gel filtration (superdex200 10 300 GL; GE Healthcare) column (using AUC Buffer; Tris25mM pH7.5, NaCl 50mM) to resolved the peak for complex. Peak fractions were analysed on 10% SDS PAGE and fractions for earlier eluting peak was pooled together, concentrated on Amicon Centricon (3kDa cutoff; Millipore) and subjected for Molecular Size determination by Analytical Ultracentrifugation Sedimentation Velocity (AUC-SV) Experiment. The AUC-SV run was performed at 42000 RPM at 20°C in Beckman XL-A ultracentrifuge. The concentrated peak fraction was diluted in AUC buffer to ∼0.8 mg/ml and loaded into standard double-sector centerpieces. The cells were scanned at 280 nm every minute and more than 300 scans were recorded. First 300 image data were processed and analysed using the program SEDFIT(Schuck, 2000) with the model of continuous c(s) distribution. The partial specific volumes of the proteins (0.742125), buffer density (1.001) and buffer viscosity (0.010142) were estimated using the program SEDNTERP. Data figures were generated using the program GUSSI.

## Cell Line and Cell Culture

U2OS cells and derivative were grown at 37 °C and 5% CO2 in DMEM (PAN Biotech) supplemented with 10% filtered Fetal Bovine Serum (FBS, Gibco), 2 mM L-glutamine (PAN Biotech), 100 U/ml penicillin and 0.1 mg/mL streptomycin (PAN Biotech, Pen/Strep mix). To obtain U2OS cells expressing the GTSE1-GFP, GTSE1^5xLID^-GFP or the CHC-GFP BAC transgenes, the corresponding BACs were transfected into U2OS cells using the Effectene kit (Qiagen) following the manufacturer’s instruction. Clonal lines expressing the BAC transgene were selected with G418 (Sigma Aldrich). The GTSE1^WT^ and GTSE1^5xLID^ clones expressing the GTSE1-GFP and GTSE1^5xLID^-GFP BAC transgenes close to endogenous level, respectively, were selected for phenotypic analysis. The mCherry ß-tubulin GTSE1^5xLID^ cells were obtained by transfecting the mCherry ß-tubulin BAC into GTSE1^5xLID^ cells and selecting a clonal line expressing both transgenes using G418 (Sigma Aldrich) and Blasticidin selection. GTSE1^KO^ cells were generated in Bendre *et al*., 2016(Bendre et al., 2016).

For the measure of PEFs and microtubule lengths in monopolar spindle, cells were treated with 10 µM STLC for 2 hours before fixation. For the cold-stability assay, cells were synchronized in G2, released for 55 minutes in normal media, transferred into ice-cold media for 17 min, and fixed immediately. Cells were synchronized in G2 32 hours post siRNA transfection by a 16 hours long 10 µM RO-3306 (Calibiochem) treatment.

## RNAi

The siRNAs 5′-GAUUCAUACAGGAGUCAAAtt-3′ (GTSE1(B3)), 5′-GAUCCAACGCAGUAAUGGUtt-3′ (MCAK), and 5’-GGUUGCUCUUGUUACGGAUtt-3’ (CHC) were routinely used to deplete GTSE1, MCAK and CHC, respectively. siRNAs GGAAUUAAAUAAUCCGGUUtt GTSE1(B1) and GUACAAAGAAGGUCACUUAtt GTSE1(T1) were used as extra-controls for GTSE1 depletion in the H3pS10 experiment. In all experiments, control cells were treated with the Silencer negative control 2 siRNA (AM4637). All siRNAs except GTSE1(T1) (Sigma) were purchased from Ambion. GTSE1 and MCAK depletion were performed using a reverse transfection approach. A transfection mix containing 2.5 µL Oligofectamine (Invitrogen) and siRNA(s) was prepared in OptiMEM (Invitrogen) and incubated 20 min at RT. Meanwhile, cells were seeded at the desired confluency into 24-well plates containing coverslips or into 8-well imaging chambers (Ibidi). The transfection mix was added to the cells and the final volume adjusted to 500 µL using pre-warmed media. Media was changed 7-8 h after transfection. Experiments were performed 48 h after transfection unless stated otherwise. GTSE1 and MCAK siRNAs were used at 80 nM and 12 nM final concentration, respectively.

CHC depletion was performed using a forward transfection approach. Cells were seeded into a 3.5 cm dish with coverslips and grown until 75% confluent. Prior to transfection, medium was exchanged for 1.8 mL OptiMEM supplemented with 100 U/ml penicillin and 0.1 mg/mL streptomycin (PAN Biotech, Pen/Strep mix). A 200 µL transfection mix containing 3µL Lipofectamine RNAi Max (Invitrogen) and the siRNA was prepared in OptiMEM (Invitrogen), and incubated 25 min at RT before addition to the cells. Experiments were performed 66 h after transfection. CHC siRNA was used at 50 nM final concentration. When necessary, cells transfected with Control or CHC siRNA for 66 h using the forward transfection approach were submitted to a second round of reverse transfection with Control, GTSE1 (100 nM), MCAK (12 nM) or CHC (100 nM) siRNAs for an extra 48 h.

## Western Blot

Samples in Laemmli buffer were run on SDS-PAGE and transferred onto a nitrocellulose membrane. Membranes were blocked with 5% milk (Carl Roth) in PBS Tween20 0.1% (SERVA electrophoresis) before incubation with the primary antibody diluted in the same blocking solution. For the mouse CHC antibody the blocking solution was changed to 5% BSA in PBS Tween20 0.1%. Secondary antibodies coupled to horseradish peroxidase were diluted in PBS Tween20 0.1% milk 5% before incubation on the membrane. For detection, membranes were incubated with ECL reagent (GE Healthcare) before imaging on the ChemiDoc^TM^ MP imaging system (BioRad) or development onto Hyperfilm (Amersham). Numeric images were obtained by scanning Hyperfilms or using the ImageLab software (BioRad) to generate tiffs from the ChemiDoc^TM^ MP files. Fiji was used to adjust levels, generate 8-bit tiffs, and crop images. Western Blot quantifications were performed on ChemiDoc^TM^ MP generated files showing no saturated pixels, and using the Volume tools of the ImageLab software. Background-corrected signals were normalized for Tubulin intensity, and expressed in percentage of the control siRNA condition. If a signal showed a negative value after background correction, the value was set to 0 to avoid problems during normalization. Samples to test RNAi efficiencies were obtained by adding hot Laemmli buffer onto cells from a 12 well plate (reverse transfection procedure scaled-up to 750 µL final volume) or by lysing cells from a 3.5 cm dish into 100 µL ice-cold Cell-lysis buffer (50 mM Na2HPO4, 150 mM NaCl, 10% glycerol, 1% Triton X-100, 1 mM EGTA, 1.5 mM MgCl2, 50 mM HEPES pH 7.2 complemented with 2X protease inhibitor mix (SERVA)).

## Immunoprecipitation and mass-spectrometry

To obtain mitotic cells, cells were synchronized with 10 µM STLC for 16 hours, harvested by mitotic shake-of, washed with PBS, incubated in media supplemented with 10 µM MG132 for 45-70 minutes at 37 °C, and harvested by centrifugation. For Aurora kinase A inhibition, 500 nM MLN8054 were applied during MG132 treatment. All cells were lysed for 15 minutes on ice in ice-cold Cell-lysis buffer (50 mM Na2HPO4, 150 mM NaCl, 10% glycerol, 1% Triton X-100, 1 mM EGTA, 1.5 mM MgCl2, 50 mM HEPES pH 7.2, 1 mM DTE) supplemented with 2X protease inhibitor mix (SERVA), and PhoStop (EASY Pack, Roche). Debris were removed by centrifugation 15 min at 18000 rcf at 4 °C. 1 mL of cell lysate at 1 mg protein / mL was used for the Immunoprecipitation. 100 µL of cell lysate were save as Input. Lysates were incubated 2 hours at 4 °C with the antibody before addition of 25 µL pre-washed Dynabeads (Invitrogen). After 4 hours incubation at 4 °C, beads were washed thrice with Cell-lysis buffer. For Western blot analysis, proteins were eluted from beads by addition of 2x hot Laemmli buffer. For mass-spectrometry analysis, IPs were performed in triplicates to obtain reliable label-free quantitative data and to perform statistical tests. Samples were reduced, alkylated and digested directly on beads with LysC/Trypsin(Hubner and Mann, 2011). Obtained peptides were desalted/concentrated on C18-reversed phase stage tips(Rappsilber et al., 2007) and then separated with a PepMap100 RSLC C18 nano-HPLC column (2, 100, 75 IDx25 cm, nanoViper, Dionex, Germany) on an UltiMate™ 3000 RSLCnano system (ThermoFisher Scientific) using a 125 min gradient from 5-60% acetonitrile with 0.1% formic acid and then directly sprayed *via* a nano-electrospray source (Nanospray Flex Ion Scource, Thermo Scientific) in a Q Exactive^TM^ Plus Hybrid Quadrupole-Orbitrap Mass Spectrometer (ThermoFisher Scientific). The Q Exactive^TM^ Plus was operated in a data-dependent mode acquiring one survey scan and subsequently ten MS/MS scans. Resulting raw files were processed with the MaxQuant software (version 1.5.2.18) searching against an uniprot human database (ref. Jan 2016) using deamidation (NQ), oxidation (M) and acetylation (N-terminus) as variable modifications and carbamidomethylation (C) as fixed modification(Cox and Mann, 2008). A false discovery rate cut off of 1% was applied at the peptide and protein level. The integrated label-free algorithm was used for relative quantification of the identified proteins(Cox et al., 2014). Quantified proteins were further analyzed with Perseus (version 1.5.1.5) (Tyanova et al., 2016). From the list of identified proteins contaminants and reverse hits were removed. To identify differences between the samples, we required proteins to be quantified in at least two out of three replicates for WT or mutant. Missing values were imputated with Perseus at the lower end of the distribution (down shift 1.8, with 0.3). The resulting 1017 proteins were used for a two-sided t-test. The result was visualized with a volcano plot (cut-off: p-value 0.01; S0 2).

## Immunofluorescence

Cells seeded onto coverslips were fixed for 10 min at RT with 4% PFA dissolved into a Pipes based buffer (50 mM Pipes, pH 7.2, 10 mM EGTA, 1 mM MgCl_2_, 0.2% Triton X-100). Membranes were permeabilized with 0.25% Triton X100 for 10 min at RT. Cells were blocked for 30 min at RT with 5% BSA (Sigma) dissolved into PBS. Primary antibodies were diluted into 5% BSA in PBS and incubated onto coverslips for 1 h at 37 °C in a wet chamber. The same procedure was applied for secondary antibodies. Coverslips were mounted using Prolong Gold antifade mounting with DAPI (Molecular Probes and Thermo Fisher Scientific). Cells were washed thrice with PBS between every step. For the assay probing the recruitment of the GTSE1 transgenes to the spindle, PBS was replaced with PBS-Tween20 0.1% (SERVA electrophoresis). For the assays measuring the spindle total tubulin fluorescence / volume, astral microtubules lengths, or microtubules lengths /PEF in monopolar spindles, cells were fixed with −20 °C methanol for 10 min. Cells were then blocked and processed as usual.

A Marianas spinning-disk confocal system (Intelligent Imaging Innovation, 3i) built on an Axio Observer Z1 microscope (Zeiss) with an ORCA-Flash 4.0 camera (Hamamatsu Photonics) was used to acquire images. Images from the microtubule cold-stability assay were acquired using a 40X 1.4 NA Plan-Apochromat oil objective (ZEISS). Images for the GTSE1 transgenes spindle localization, TACC3 localization, and astral microtubules length measurement following CHC depletion were taken using a 100× 1.4 NA Plan-Apochromat oil objective (ZEISS). Other images were taken with a 63× 1.4 NA Plan-Apochromat oil objective (ZEISS).

## Live cell imaging

Culture media was exchanged for CO2-independent visualization media (Gibco) 5 hours prior imaging. To visualize DNA or microtubules, 1 hour prior to imaging cells were treated with 1 µM Verapamil and 1 µM siRDNA or 1 µM siRTubulin (Spirochrome), respectively. To measure the time between NEB and anaphase onset, cells were imaged at 37 °C overnight. Images of U2OS, GTSE1^WT^ and GTSE1^5xLID^ cells were acquired at 2 minutes intervals in series of 2 µm spacing z-sections. Images of mCherry ß-tubulin GTSE1^5xLID^ cells were acquired at 3 minutes intervals in series of 2.5 µm spacing z-sections. For the localization of the GTSE1 transgenes in GTSE1^WT^ and GTSE1^5xLID^ cells, images were acquired in series of 0.2 µm spacing z-sections. For the chromosome congression movies, images were acquired at 1.5 minutes intervals in series of 1 µm spacing z-sections. All Live Cell imaging was performed on a DeltaVision imaging system (GE Healthcare) using a an sCMOS camera (PCO Edge 5.5) and a 40x 1.3NA UPLFLN 40XO objective (Olympus) or a 60x 1.42NA Plan Apo N UIS2 objective (Olympus). When required, images were deconvolved using the SoftWoRx software 6.1.l and/or z-projected (Max intensity) using the SoftWoRx software 6.1.l or Fiji 1.0.

## Microscopy image analysis and quantifications

Quantifications were performed using the Imaris 7.6.4 software (Bitplane). Images were opened and analysed in 3D in the ‘Surpass’ mode. The ‘Surface’ function was used on the α-tubulin channel to isolate spindles, half-spindles, and cold-resistant microtubules and on the H3pS10 channel to isolate the region of the DNA showing H3pS10 signal. The ‘Spot’ function was used on the EB1, CREST, MAD1 and PCNT channels to define EB1 comets, kinetochores, MAD1 foci and spindle poles, respectively. To measure backgrounds, spots were manually placed either in the cytoplasm or outside of the cells using the ‘Spot’ function. The 3D coordinates *(x,y,z)*, volume, prolate ellipticity, or fluorescence intensities of the objects defined with the ‘Spot’ and ‘Surface’ functions were retrieved through the ‘Statistics’ option. The recruitment of a protein of interest (POI) to the spindle or half spindle was calculated as the 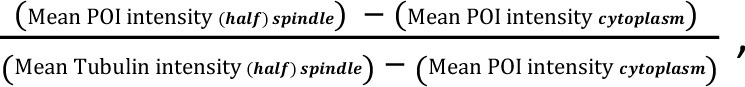 where the Mean intensity of a POI is the mean fluorescence intensity of the corresponding channel in the surface / dot of interest. For TACC3 and CHC-GFP the Mean tubulin intensity was not corrected for cytosolic background. The total spindle-Tubulin and H3pS10 fluorescence were calculated as the total fluorescence intensity of the corresponding channel in the object of interest (Sum Intensity measurement in Imaris) corrected for background. In this case, the background was measured in a spot of fixed volume, and the background for an object of the same volume as the object of interest was calculated: 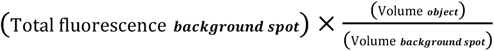. microtubules, the cold-resistant microtubules of a cell were defined as one or multiple object(s) using the ‘Surface’ function. The tubulin fluorescence intensity of each object was defined as: (Mean tubulin intensity *object* – Mean tubulin intensity *cytoplasm*) × (Volume *object*). If more than one object were present in a cell, the tubulin fluorescence of all the objects were summed to get the total tubulin fluorescence of cold-resistant microtubules. For the quantification of the percentage of misaligned chromosomes with MAD1 positive kinetochores, a kinetochore was defined as 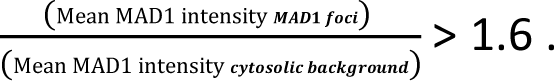 Because the MAD1 foci do not fully overlap with the CREST signal, MAD1 foci were defined manually using the ‘Spot’ function. Based on the localization of their kinetochores, chromosomes were classified in three categories: 1) outside the inner-spindle, 2) inside or contacting the inner-spindle, and 3) a category grouping chromosome with ambiguous localization or associated with a detached pole.

Distances between two points were calculated using the 3D coordinates (*x,y,z*) of these points and the following formula 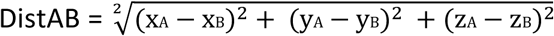. Distances between EB1 comets and the closest spindle pole or the spindle monopole were used as a proxy for microtubule lengths. PEFs were estimated by calculating the distances between kinetochores and the monopole. The monopole was defined as the middle point between the two centrosomes (PCNT dots) using the formula 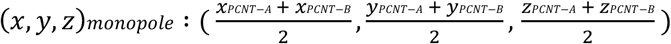. The method to measure astral microtubule lengths was adapted from (Stout et al., 2011). Spindle poles were defined either manually using the ‘Spot’ function and the tubulin and EB1 signal as a reference or using the ‘Spot’ function on the PCNT staining. Distances between EB1 comets and their closet spindle pole were calculated. The angle between the line formed by a comet and its closet pole and the line formed by the two spindle poles was calculated using the cosine law. An EB1 comet was considered as belonging to an astral microtubule when this angle was superior to 60°. The number of astral microtubules per cell and the average astral microtubule length per cell were then calculated. All analysis were performed in Microsoft Excel or R. Some data were exported from IMARIS in Excel files and imported in R using the XLConnect package.

## Statistical analysis and plot generation

All statistical analysis, except the Mass spectrometry analysis (*see corresponding section*), were performed in R 3.4.3. Usually, experiments were analyzed as follow. The normality and homoscedasticity of the data were tested using a Shapiro and a Bartlett test, respectively. If the normality and homoscedasticity of the data were verified, differences were evaluated using an ANOVA followed by a Tukey. If normality or homoscedasticity were not verified, differences were assessed using a Kruskal Wallis test followed by direct comparisons using a Wilcoxon test. Occasionally, a Student test was used for small data set, typically n=3 (See Table S1 for detailed statistical analysis). The ggplot package was used to generate plots that were exported as pdf files. The generated pdf files were imported and edited in Illustrator CS5.1.

## Evolutionary analysis

To trace the origin of *GTSE1* among vertebrates, we first inspected a multiple alignment between the human and many other vertebrate genomes {PMID 28645144}. This revealed alignments to the most vertebrates including the spotted gar and elephant shark, but not the two lamprey species (Petromyzon marinus and Lethenteron camtschaticum) that represent the most basal vertebrate lineage. For the elephant shark, we inspected aligned ESTs and protein sequences, which revealed that the genome contains a full length *GTSE1* gene that aligns over the entire length to GTSE1 of other vertebrates (Fig. S3). Furthermore, the elephant shark GTSE1 is located in a genomic region that contains the same gene order (*ALG10*, *TPRKB*, *GTSE1*, *TRMU*, *CELSR1*) as in the spotted gar, suggesting that this is the ancestral gene order. In mammals, gene order upstream of GTSE1 was changed by genomic rearrangements. BLAST also identified a full length GTSE1 protein in the whale shark (Fig. 2 B, Fig. S3). To further probe the presence of *GTSE1* in lamprey, we queried Lamprey proteins provided by SIMRbase (https://genomes.stowers.org/organism/Petromyzon/marinus) with blastp, which revealed that the lamprey possesses a *GTSE1* gene (protein identifier PMZ_0002060-RA) that contains the N-terminal and the C-terminal domains. Next, we used the gar and elephant shark GTSE1 protein sequence to perform sensitive searches against the non-vertebrate portion of the NCBI non-redundant protein sequence (NR) database using PSI-BLAST with word size 2, gap existence/extension costs of 10 and 3, and the BLOSUM45 matrix. We used the human TACC3 protein to run PSI-BLAST searches against the NCBI NR database and blastp searches against SIMRbase to retrieve homologs in lamprey, ciona, sea urchin and Drosophila. Multiple alignments of GTSE1 and TACC3 were created with MUSCLE v3.8.31 {PMID 15034147} and default parameters.

---

**Figure S1:**
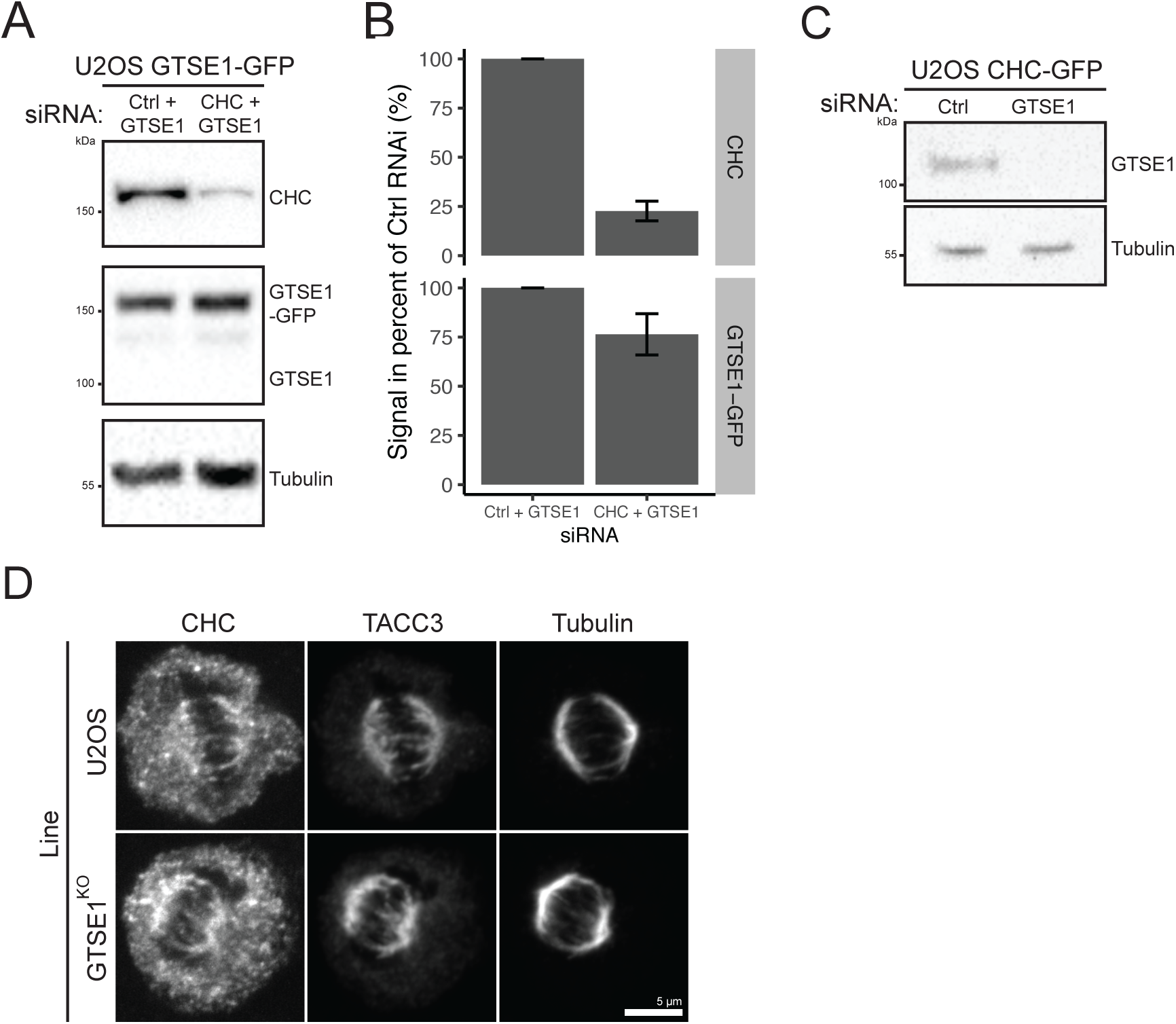
Related to Figure 1, RNAi depletion efficiencies and localization of CHC and TACC3 in U2OS cells knocked out for GTSE1. **(A)** Immunoblot on the cell lysate of U2OS cells expressing an siRNA resistant GTSE1-GFP BAC transgene and transfected with control (Ctrl) or CHC siRNA for 66 hours. Cells were concomitantly transfected with GTSE1 siRNA to deplete endogenous GTSE1. Immunoblots are with anti-GTSE1, anti-CHC, or anti-α-tubulin antibodies. Anti-CHC blot was performed on same samples run on a different gel. **(B)** Quantification of CHC and GTSE1-GFP on immunoblot of three siRNA experiments performed as in (A). GTSE1-GFP and CHC levels were normalized to α-tubulin and presented in percent of the Ctrl siRNA. Mean percentages (+/-SE) are presented. (N=3 exp.). **(C)** Immunoblot on the cell lysate of U2OS cells stably expressing a CHC-GFP BAC transgene and transfected with Ctrl or GTSE1 siRNA. The immunoblot is with anti-GTSE1 and anti-α-tubulin antibodies. **(D)** Immunofluorescence images of U2OS cells and U2OS cells knocked-out for GTSE1 (GTSE1^KO^), stained with antibodies against CHC, TACC3 and α-tubulin. Scale bars: 5 µm. Numeric data in Table S1.

**Figure S2:**
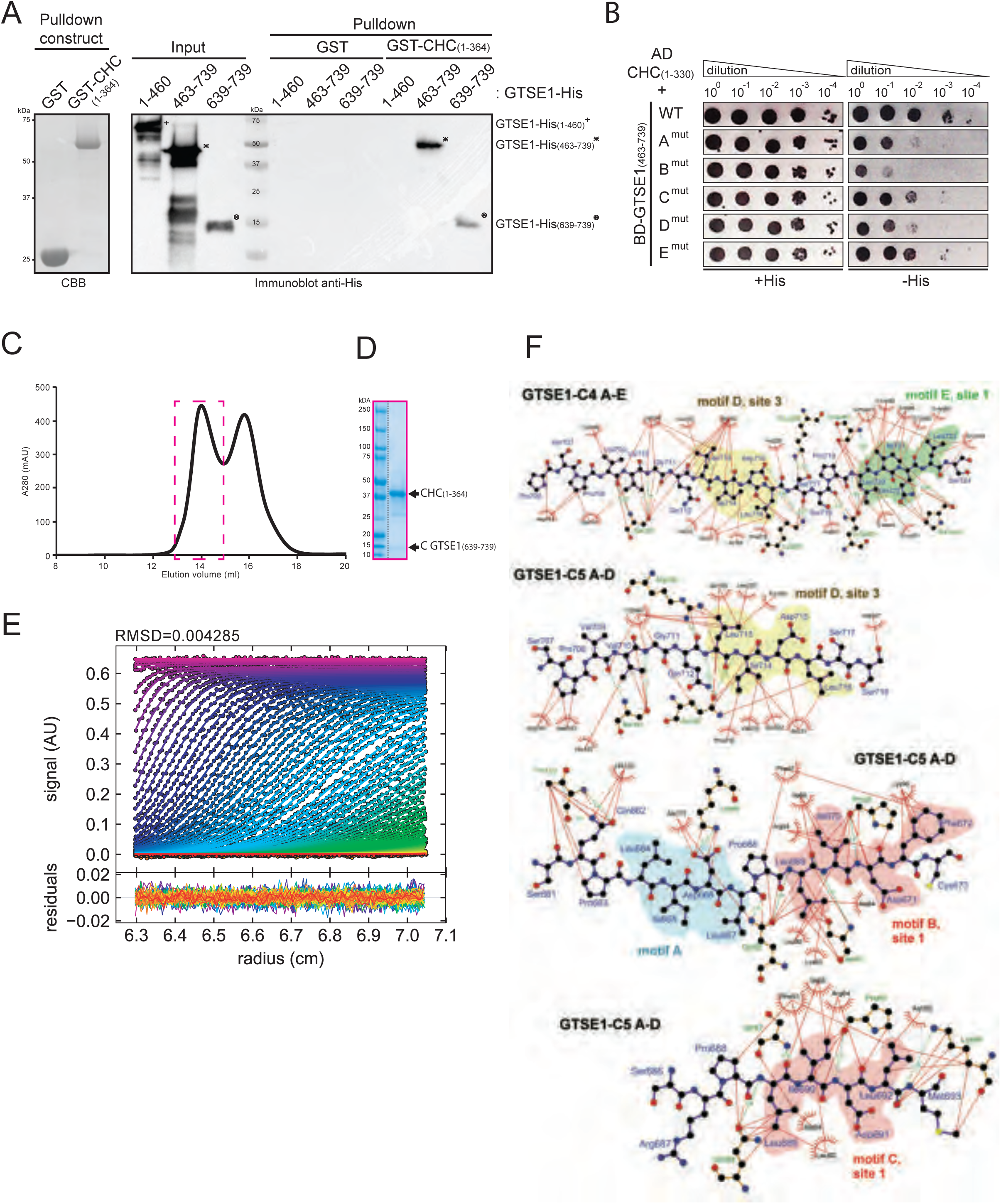
Related to Figure 2, Clathrin box motifs on the GTSE1 C-terminus interact with adaptor binding sites on the CHC N-terminal domain. **(A)** GST-pulldown analysis of the interaction between GST-CHC(1-364) and GTSE1 fragments fused to a His-tag (GTSE1-His). Numbers indicate a.a. present in the fragments. Immunoblot is with anti-His antibody. **(B)** Yeast-two-hybrid analysis of the interaction between CHC(1-330) fused to activation domain (AD) and different GTSE1(463-739) fragments fused to the DNA binding domain (BD). Letters A to E indicate mutated motifs in GTSE1 (see Fig. 2B for sequences). Interaction was visualized by yeast growth on -His selection plates. **(C)** Gel filtration profile (superdex200 10 300 GL column) of CHC(1-364) in complex with GTSE1(639-739) fused to 6 His tag. The indicated (pink box) peak protein fraction was run on SDS-PAGE (Coomassie blue stained) **(D)**. Fractions containing CHC(1-364) in complex with GTSE1(639-739)6xHis were pooled and submitted to AUC-SV. **(E)** AUC-SV data best curve fitting results with RMSD value in top. **(F)** Ligplot showing the amino-acids involved in the interactions between CBMs on GTSE1 and Site 1 and 3 on the CHC-NTD. The Ligplot were designed based on the crystal structures of GTSE1-C4(661-726) (upper panel, PDB ID: 6QNN) and GTSE1-C5(653-719) (lower panel, PDB ID: 6QNP) in complex with CHC(1-364). Site 3 is occupied by motif D in both crystals while Site 1 is occupied by motif E in the GTSE1-C4 crystal and by motif B or motif C in the GTSE1-C5 crystal. Amino-acids constituting CBMs are highlighted.

**Figure S3:**
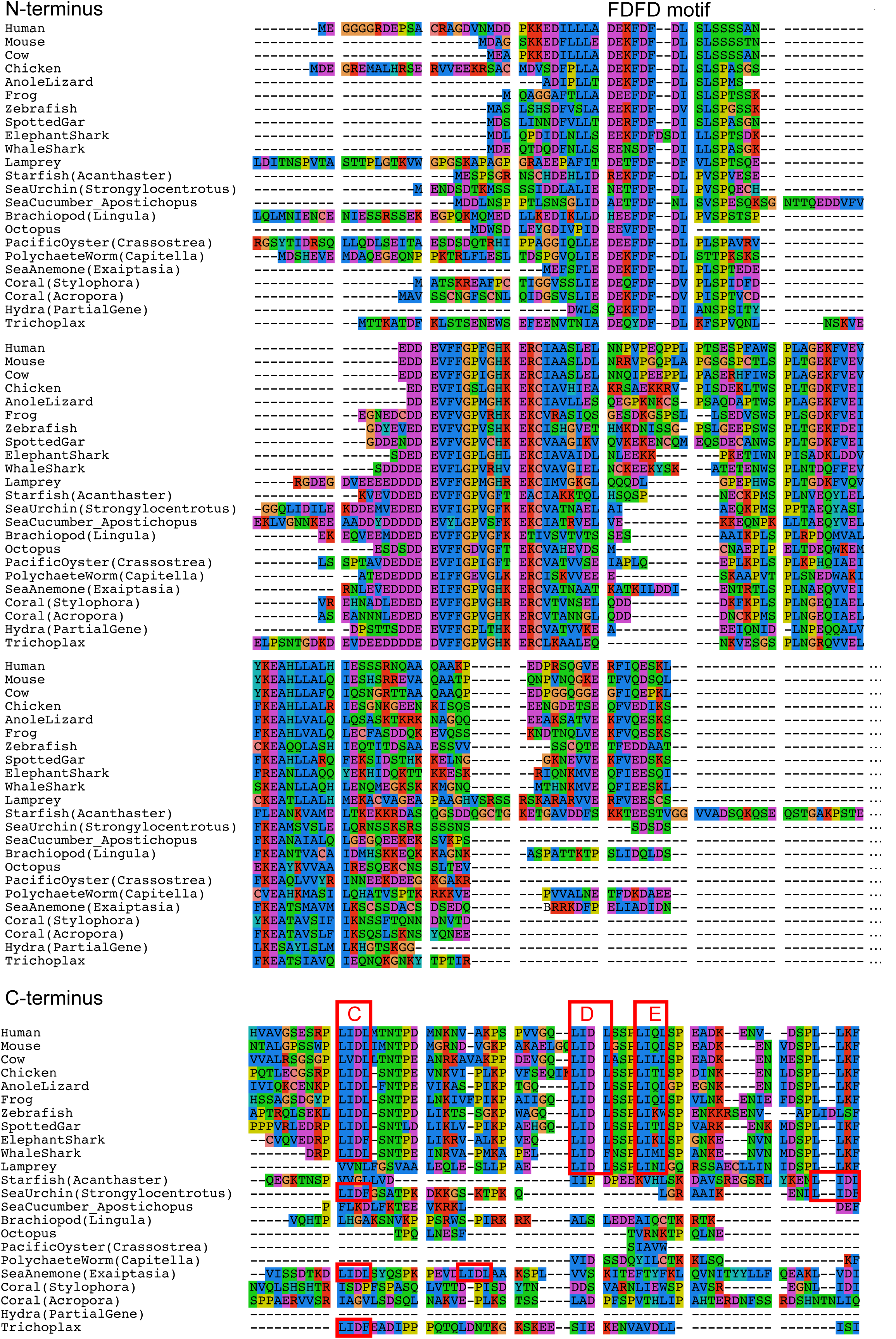
Related to Figure 2, Alignment of the full-length GTSE1. MUSCLE alignment of the N-(amino-acids 1-143 from human GTSE1) and C-(amino-acids 679-739 from human GTSE1) termini of potential GTSE1 homologs. Alignment was performed on full-length genes from the indicated organisms. A conserved N-terminal FDFD motif is labelled. Three putative clathrin binding box motifs (C to E) are indicated in red. Notice some non-vertebrates also have putative clathrin binding box motifs in the C-terminal region (red boxes).

**Figure S4:**
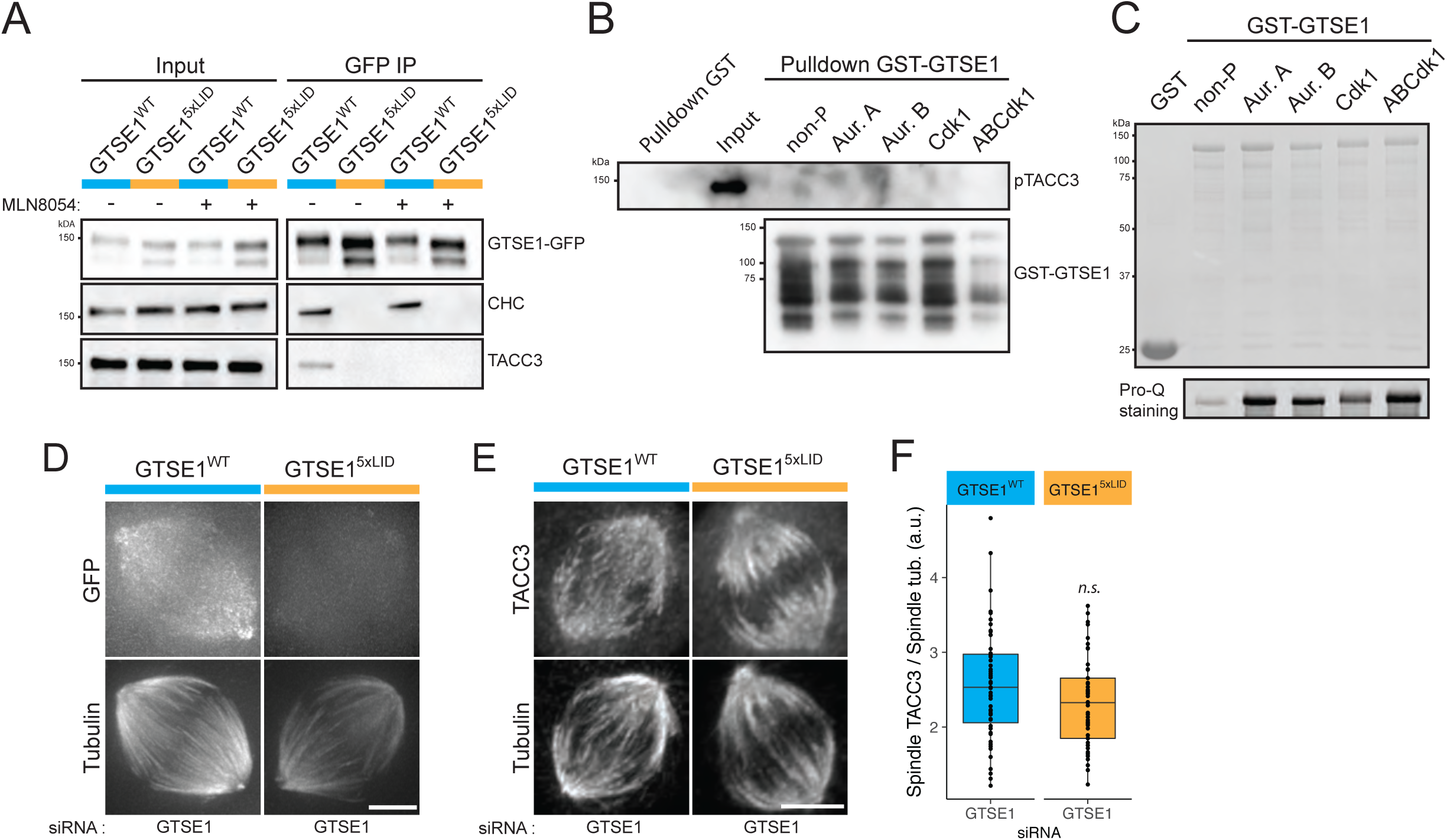
Related to Figure 3, The CHC-GTSE1 interaction recruits GTSE1 to the CHC/TACC3 complex and the spindle but does not control TACC3 spindle localization. **(A)** Immunoblot on mitotic lysates of GTSE1^WT^ and GTSE1^5xLID^ cells (Input) and immunoprecipitations of the GFP transgene (GFP IP). MLN8054 was used to inhibit Aurora A. Immunoblot are with anti-GFP, anti-CHC or anti-TACC3. **(B)** GST and GST-GTSE1 pulldowns of purified TACC3 phosphorylated by Aurora Kinase A. GST-GTSE1 was left unphosphorylated or was phosphorylated by Aurora A (Aur. A), Aurora B (Aur.B), Cdk1 or Aurora A, Aurora B and Cdk1 (ABCdk1). Phosphorylated TACC3 and GTSE1 were visualized by immunoblot against pTACC3(S558) and GTSE1, respectively. **(C)** SDS-PAGE stained with Coomassie blue of GST, unphosphorylated GST-GTSE1 and GST-GTSE1 phosphorylated by Aurora A (Aur. A), Aurora B (Aur.B), Cdk1 or Aurora A, Aurora B and Cdk1 (ABCdk1). Presence of phosphorylated residues was confirmed by Pro-Q staining. **(D)** Still images from live-cell imaging on GTSE1^WT^ and GTSE1^5xLID^ cells transfected with GTSE1 siRNA to deplete the endogenous GTSE1. Cells were treated with siRTubulin prior to imaging to visualize microtubules. (E) Immunofluorescence images showing the localization of TACC3 to the spindle in GTSE1^WT^ and GTSE1^5xLID^ cells transfected with the GTSE1 siRNA. TACC3 and α-tubulin staining are shown. **(F)** Quantification of TACC3 on the spindle of cells from (E). Mean TACC3 fluorescence intensity on the spindle was corrected for background and normalized to α-tubulin (n>50 cells per condition over N= 3 experiments pooled for analysis and representation, *p-value* from ANOVA). All scale bars: 5 µm. Numeric data in Table S1.

**Figure S5:**
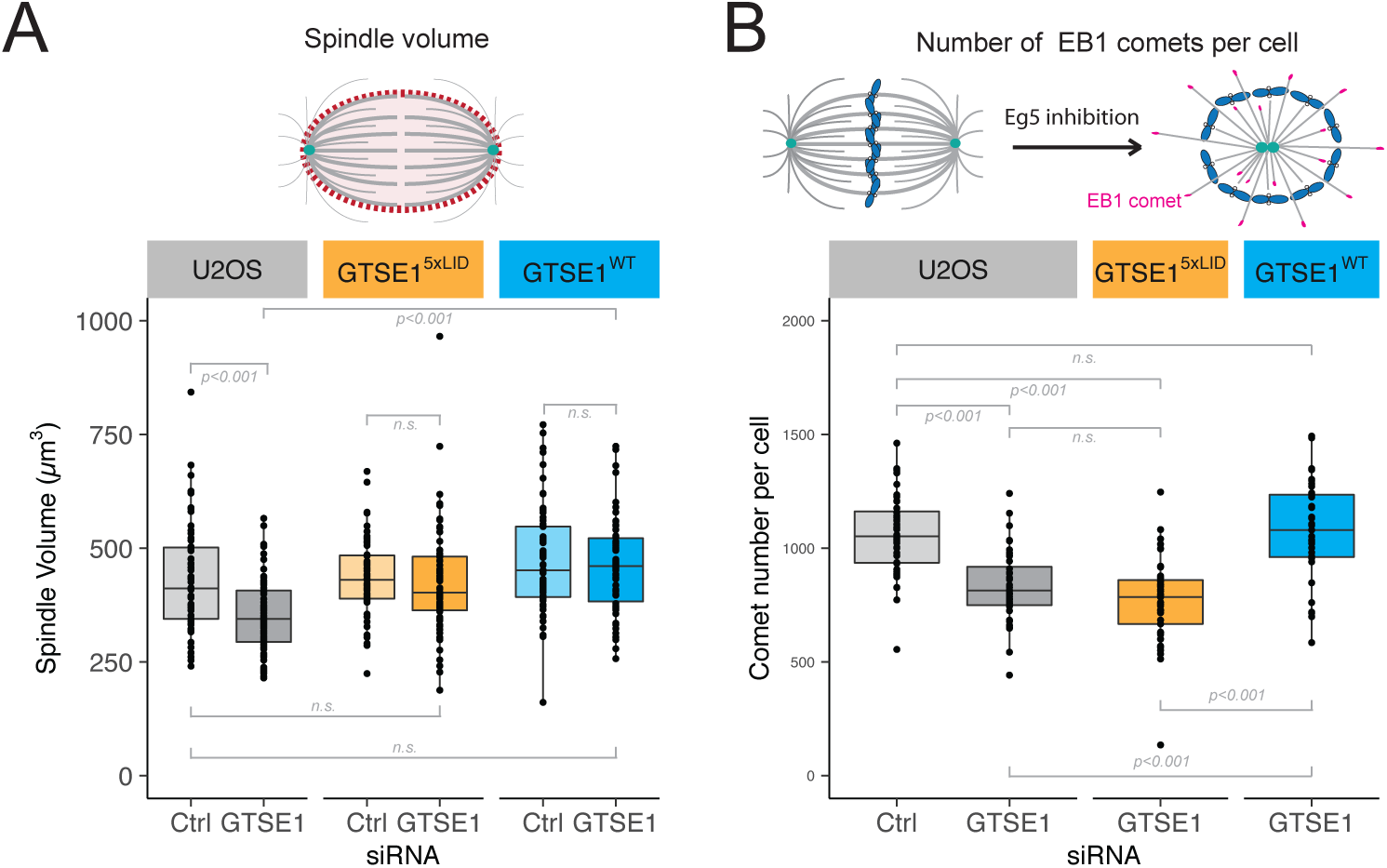
Related to Figure 4, Further analysis of the effect of disrupting the CHC-GTSE1 interaction on mitotic spindles. Quantification of the spindle volume **(A)** in U2OS, GTSE1^WT^ and GTSE1^5xLID^ cells, transfected with Control (Ctrl) or GTSE1 siRNA. Quantifications were performed on cells from Fig. 4 A using 3D objects representing the spindles without astral microtubules. (n>60 cells per condition, N=2 experiments pooled for representation and analysis, *p-values* from Wilcoxon tests in (B)) **(B)** Number of EB1 comets in U2OS, GTSE1^WT^ and GTSE1^5xLID^ cells, transfected with Ctrl or GTSE1 siRNA and treated with STLC to form monopolar spindles. Quantifications were performed on cells from Fig. 4 F using 3D objects reconstruction (n>36 cells per condition, N=3 experiments pooled for analysis and representation, *p-values* from Wilcoxon test). Numeric data in Table S1.

**Figure S6:**
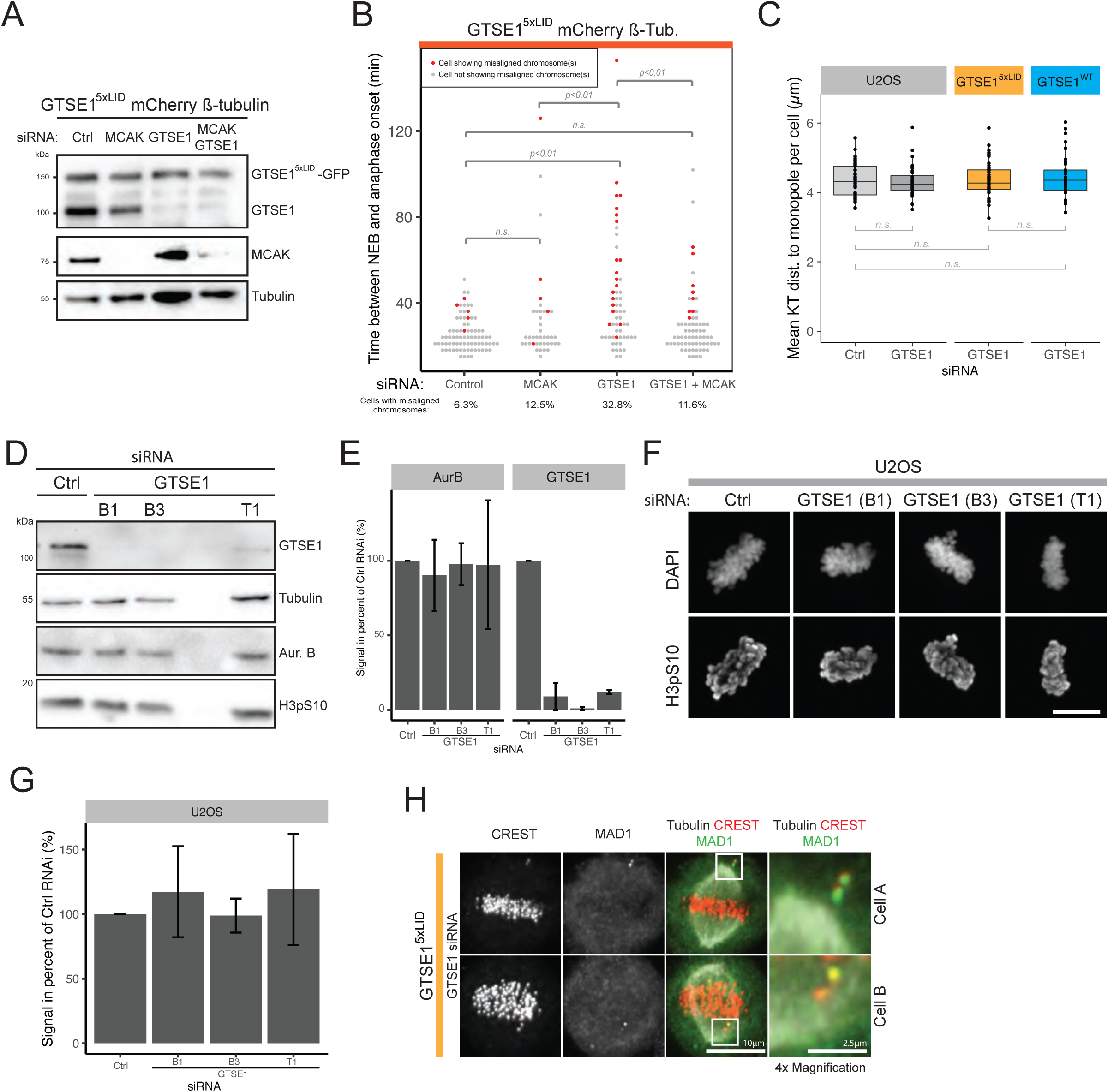
Related to Figure 5, Analysis of chromosome congression in GTSE1^5xLID^ cells and quantification of Aurora B kinase abundance and activity after GTSE1 depletion. **(A)** Immunoblot on asynchronous cell lysates of GTSE1^5xLID^ cells expressing a mCherry ß-tubulin BAC transgene and transfected with Control (Ctrl), GTSE1, MCAK or GTSE1+ MCAK siRNA(s). Immunoblots are with anti-GTSE1, anti-α-tubulin or anti-MCAK. **(B)** Time from nuclear envelope breakdown (NEB) to anaphase onset in cells from (A). Cells showing misaligned chromosome(s) are indicated in red. The percentage of cells with misaligned chromosomes is indicated. (n>39 cells, N=1, *p-value* from Wilcoxon test). **(C)** PEF assay. Mean kinetochore (CREST) to monopole (PCNT) distance per cell following 3D reconstruction from experiment shown in Fig. 4 F are presented (n>36 cells per condition, N=3 experiments pooled for analysis and representation, *p-values* from Wilcoxon test). **(D)** Immunoblot on mitotic lysates of U2OS cells transfected with Ctrl, GTSE1(B1), GTSE1(B3) or GTSE1(T1) siRNA. Cells were synchronized in mitosis with Nocodazole. Phosphorylation of H3 Serine 10 (H3pS10) is used as a marker of Aurora B activity. Immunoblots are with anti-GTSE1, anti-Aurora B, anti-H3pS10, or anti-α-tubulin antibodies. **(E)** Quantification of GTSE1 and Aurora Kinase B levels in immunoblots of U2OS cells transfected with Ctrl or GTSE1 siRNA(s). Levels were first normalized to α-tubulin level and then to Ctrl siRNA. Cells were synchronized in mitosis with Nocodazole. The mean +/-SD is presented, N=2. **(F)** Immunofluorescence images of metaphase-like U2OS cells transfected with Ctrl, GTSE1(B1), GTSE1(B3) or GTSE1(T1) siRNA. DNA and H3pS10 staining are shown. Scale bar: 10 µm. **(G)** Quantification of the H3pS10 total fluorescence intensity in cells from (F). The total H3pS10 fluorescence intensity per cell was measured from 3D reconstructions and corrected for background. Within each experiment, the average total H3pS10 fluorescence intensity per condition was calculated and normalized to the corresponding Ctrl siRNA. The mean of these normalized values (+/-SD) over N=2 experiments is presented (n>36 cells per condition). **(H)** Immunofluorescence images of GTSE1^5xLID^ cells transfected with GTSE1 siRNA to deplete the endogenous GTSE1 and showing a misaligned chromosome outside (upper lane) or within the spindle (lower lane). Tubulin, CREST and MAD1 staining are shown. 4 x magnification of the misaligned chromosomes are presented. Positions of the magnified area are indicated. Numeric data in Table S1.

**Fig. S7:**
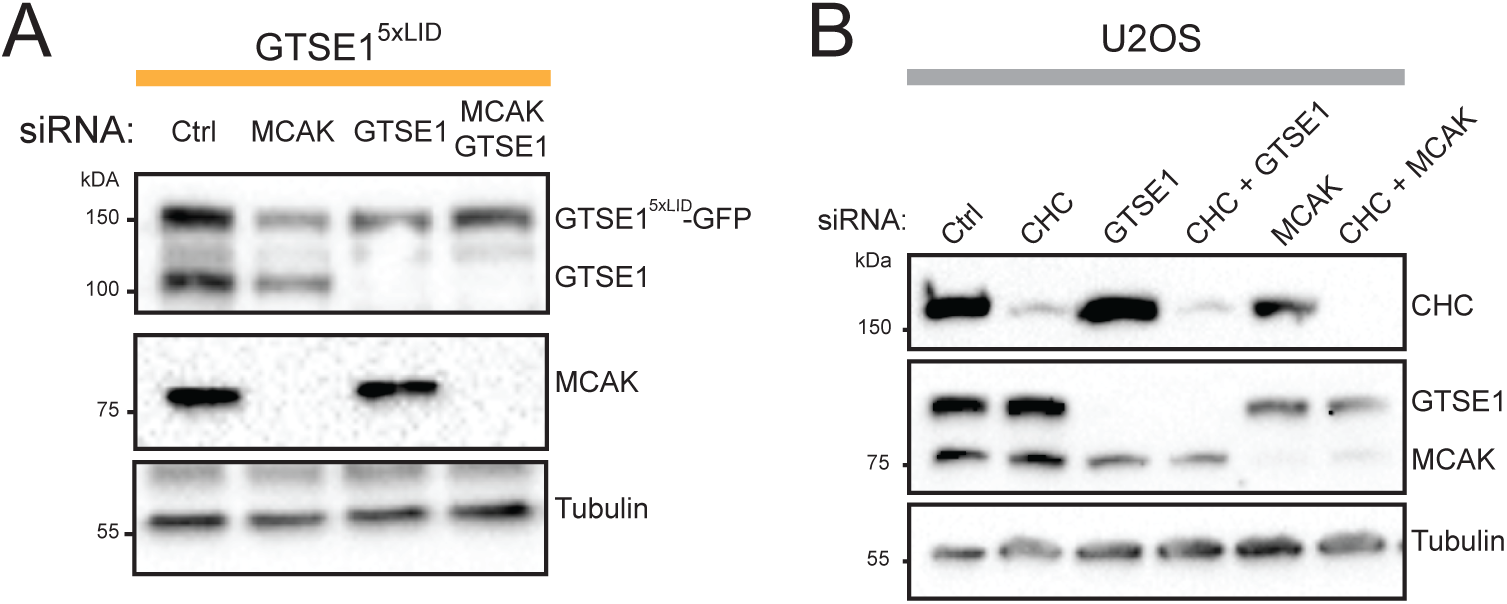
Related to Figure 6, efficiencies of RNAi procedures. **(A)** Immunoblot on asynchronous cell lysates of GTSE1^5xLID^ cells transfected with Control (Ctrl), GTSE1, MCAK or GTSE1+ MCAK siRNA(s). Immunoblots are with anti-GTSE1, anti-α-tubulin or anti-MCAK. **(B)** Immunoblot on asynchronous cell lysates of U2OS cells transfected with Ctrl, CHC, GTSE1, MCAK, CHC + GTSE1 or CHC + MCAK siRNA(s). Immunoblots are with anti-CHC, anti-GTSE1, anti-MCAK, or anti-α-tubulin.

**Fig. S8:**
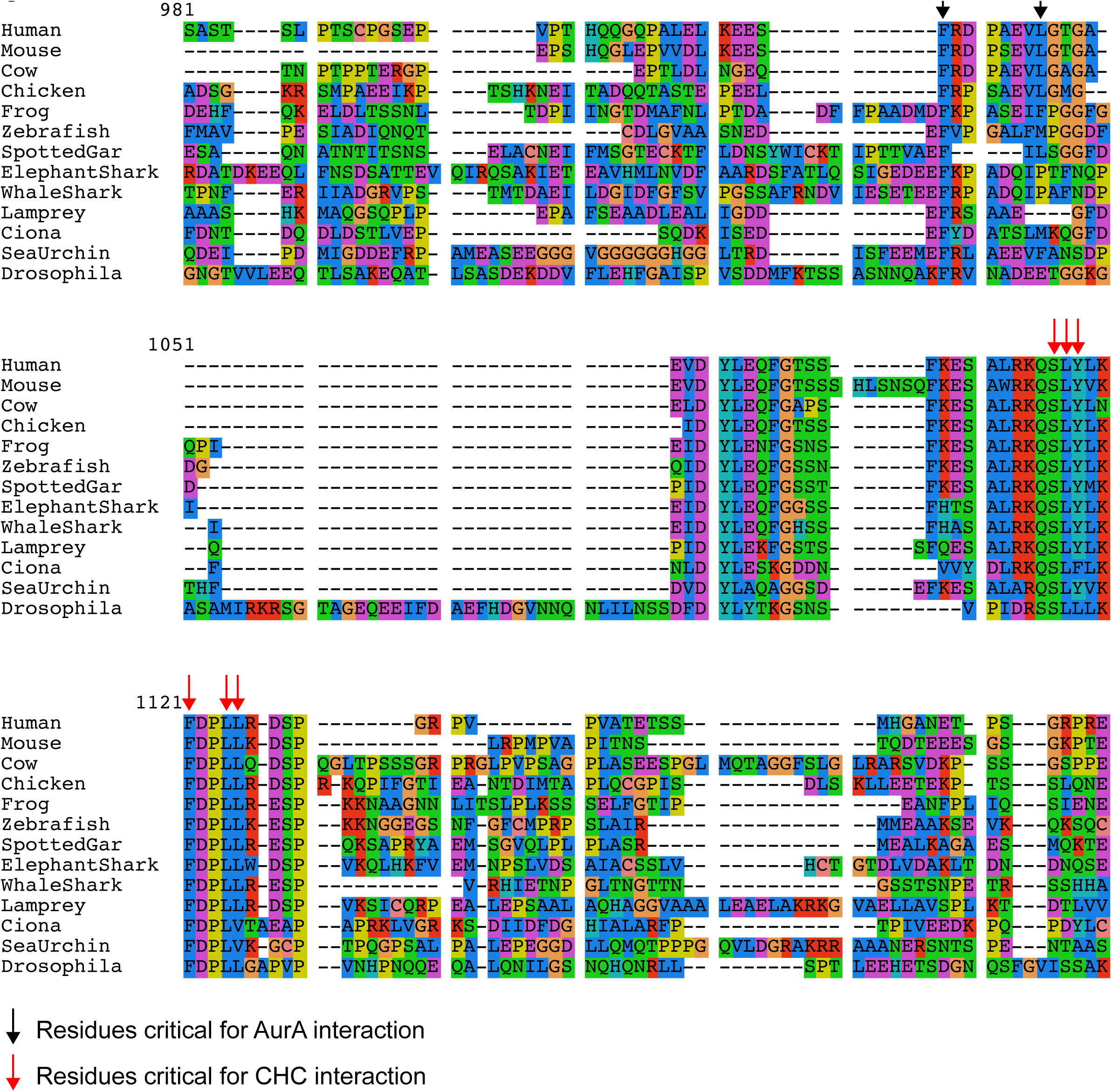
Related to Figure 2, MUSCLE alignment of the region of TACC3 that interacts with Aurora A and CHC. Alignment was performed on full-length TACC3 homologs from the indicated organisms using MUSCLE. Only the region encompassing the described interaction sites between TACC3 and Aurora A / CHC is presented. Human residues S558, L559, Y560, F563, L566 and L567 (important for CHC interaction) and F525 L532 (important for Aurora A interaction) are designated by red and black arrows, respectively. These residues appear conserved in the three non-vertebrate species carefully analyzed (sea squirt, sea urchin, and fruit fly).

**Table S1: Numeric data used to generate the plots presented in this study.**

**Table S2: Related to Figure 2, X-ray data collection and refinement statistics for the CHC(1-364) / GTSE1 peptide complexes and GTSE1 residues visible in the electron densities.** The CHC NTD (1-364) was crystallised in complex with GTSE1-C4 (containing putative CBMs A to E, PDB ID: 6QNN) or with GTSE1-C5 (containing putative CBMs A to D, PDB ID: 6QNP). **(A)** X-ray data collection and refinement statistics. **(B)** GTSE1 residues present in sites 1 and 3 of the CHC NTD. The crystal of the CHC(1-364)/GTSE1-C4 complex contained 1 CHC monomer per asymmetric unit (chain A), while the crystal of the CHC(1-364)/GTSE1-C5 complex contained 4 monomers per asymmetric units (chain A, B, C and D). Residues of the GTSE1 peptides that are visible in the electron density are presented, with GTSE1 motif A coloured in red, motif B in green, motif C in orange, motif D in blue, and motif E in magenta.

**Table S3: Related to Figure 3, Mass spectrometry data for the immunoprecipitation of the GTSE1-GFP and GTSE1^5xLID^-GFP transgenes in mitotic lysates.**

**Supplementary video 1: Related to Figure 5, Chromosome congression in GTSE1^5xLID^ cells expressing mCherry ß-tubulin and transfected with GTSE1 siRNA.** Movie based on the live cell imaging of GTSE1^5xLID^ cells expressing a mCherry ß-tubulin BAC transgene and transfected with GTSE1 siRNA. Cells were treated with siRDNA to visualize DNA. Movie from top and side view are based on a 3D reconstruction in IMARIS. Scale bar: 10 µm.

## Supplementary methods and information: CHC/GTSE1 complexes structural analyses

### Crystallization

The complex of CHC(1-364) with the GTSE1 peptide C4 (comprising “LIDL”-motifs A-E) was concentrated to 32 mg/ml after the gel filtration in 20 mM TRIS pH 7.5, 50 mM NaCl. The sequence of this construct is *GPLG*-^661^SQPLIDL…QLSPE^726^, where *GPLGS* is a cloning artifact from the PreScission site. Crystals were grown at 20° with a reservoir solution of 20% PEG 6000, 0.1M TRIS pH 8.0 and frozen using PEG300 as cryoprotectant.

The complex of clathrin with GTSE1 peptide C5 (encompassing motifs A-D *GPLGS*-^653^LAV…SSP^719^) was purified via gel filtration in 20 mM TRIS pH 7.0, 50 mM NaCl and concentrated to 30 mg/ml. Crystals grew at 20° with a reservoir of 12% PEG4000, 0.1M sodium acetate, 0.1M HEPES pH 7.5 and were frozen using PEG300 as cryoprotectant.

### Structure solution

Data was collected at 100K using a Pilatus 6M detector at the X10SA beamline of the SLS in Villigen, Switzerland. All data sets (Table S2A) were integrated and scaled using XDS and XSCALE(Kabsch, 1993). The structures were solved via molecular replacement with PHASER(Collaborative Computational Project, Number 4, 1994) in space groups P43 with a unit cell of 52 52 154 90 90 90 and one molecule per asymmetric unit for the complex with GTSE1-C4 (A-E), or in space group P21 and cell constants 95.1 90.2 100.2 90.0 110.1 90.0 with 4 molecules per asymmetric unit for the complex with GTSE1-C5 (A-D), using the clathrin moiety of 1UTC or 5M5T as template. The P21 dataset exhibits translational pseudosymmetry corresponding to an approximate doubling of the unit cell in direction of the x axis. This configuration in the crystal leads to every second reflection in this direction being relatively weak, but the processing in the true cell resulted in much clearer electron density compared to refinement trials in the smaller unit cell. Significant differences in the densities and in the positions of the molecules in the larger cell confirmed that refinement in the smaller cell would not be correct.

Example Fo-Fc densities of the A-D and A-E peptides are shown in Fig Struct S1 and Struct S2 (at the end of this document) Refinement with PHENIX (Adams et al., 2010) and building with COOT(Emsley et al., 2010) resulted in models with good Ramachandran geometry and reasonable R_work_/R_free_ values (Table S2A). GTSE1 residues identified in the electron density are listed in Table S2B.

**For the P43 structure with GTSE1-C4 (motifs A-E),** data to 2.03Å was used in spite of the relatively high Rsym values since this improved the convergence and quality of the refinement, resulting in R_work_/R_free_ values of 23.6% and 28.2%, respectively, and 96.8% residues in favored regions of the Ramachandran plot with no outliers. The sequence of the bound peptide could be assigned unambiguously to motif D of GTSE1 (“LIDL”) in the “site 3” of the clathrin molecule and to the motif E (“LIQL”) that is bound to site 1 of a neighboring, symmetry-related clathrin molecule. Details of the interaction are shown in FigS2 C. Since pull-downs with the GTSE1 peptide “C3” (residues 693-739) which starts after motif C and contains only motifs D and E still show binding to clathrin with a mutated motif E, but no binding upon mutation of the “LIDL” sequence of motif D to either AAAL, LAAA or AADA (Fig 2G), the interaction of motif D with site 3 of clathrin seems to be the dominant one. In the crystal, the GTSE1 peptide shows continuous density for the linker residues ^717^SSP^719^ between motifs D and E. This is far too short for a simultaneous binding to site 1 and site 3 of the same clathrin molecule. The GTSE1 residues preceding motif D show well-defined electron density starting from proline 706. Although ^709^VVGQ^712^ are contacting a symmetry related molecule, the three N-terminal residues ^706^PSP^708^ are not influenced by crystal packing. An amphiphysin peptide (ETLLDLDFLE) in complex with clathrin (PDB ID 5M5T) (Muenzner et al., 2016) shows a quite similar conformation to GTSE1 in both sites 1 and 3, but with fewer visible residues in site 3 (ET**LLDL** compared to PSPVVGQ**LIDL** of GTSE1, equivalent residues are in bold). In contrast, Arrestin2L binds very differently in site 3 (PDB ID 3GD1; (Kang et al., 2009)), probably due to the fact that the binding region (^332^GG**LL**GD**L**AS^340^, “splice loop”, interacting leucines in bold) is part of a much larger domain, thus restraining its conformation.

Different from the clathrin-amphiphysin complex (5M5T), Site 4 is occupied by a symmetry related molecule in our structure. Site 2 in the central opening of the clathrin beta propeller shows a relatively strong, forked density that looks different from water or a PEG molecule, but can not be assigned reliably, it could perhaps be an arginine side chain of another region of the GTSE1 peptide.

Since no higher oligomers are observed upon analysis of the complex by AUC (Fig. 2 D;Fig. S2, C-E) or SEC-MALS (data not shown), it is likely that the configuration of GTSE1 motifs D and E bridging two clathrin molecules in the crystals is serendipitous. Indeed, the presence of the peptide leads to a change of the crystal form, since the apo clathrin crystallizes in space group C2 in similar conditions (20% PEG 3350, 0.2M sodium formate, data not shown).

**For the P21 structure with GTSE1-C5 (motifs A-D)**, the initial tight NCS constraints were gradually relaxed, thus improving the R_work_/R_free_ values to 24.1% and 29.1%, respectively. 96.4% of the residues are in favored regions of the Ramachandran plot with no outliers. In all four clathrin monomers in the asymmetric unit, GTSE1-peptide density is visible in sites 1 and 3, with sites 3 having the strongest density. It could be clearly seen that motif D is bound in all four sites 3 of the four clathrin molecules. Similar to the GTSE1-C4 complex structure described above, the N-terminal part of GTSE1 motif D (^707^SPVVGQ^712^) is clearly visible. The peptides bound to site 3 of the clathrin monomers A and B do not participate in any crystal contacts, so it is evident that these positions are not a crystal packing artifact. Thus, it is likely that the PVVG motif enhances the affinity of motif D to site 3, as confirmed by the loss of affinity upon mutation of this motif (Fig S2E).

The GTSE1 peptide densities in sites 3 are not connected to the (weaker) densities in sites 1, so site 1 could harbor any motif of GTSE1, and it is not evident if the two peptides in site 1 and site 3 belong to the same GTSE1 molecule (either bound to the same clathrin or connecting two symmetry-related clathrin molecules). Although the densities for the peptides bound in sites 1 were a bit difficult to interpret, there are clear differences between the densities in sites 1 of monomers A and B compared to the sites 1 of monomers C and D. Details of the interaction are shown as a LigPlot(Laskowski and Swindells, 2011) in FigS2 C. For monomers C and D, the density in site 1 was assigned to motif B of GTSE1 based on the larger density for the phenylalanine in the sequence ^666^DLPLID**F**^672^, as well as on the side chain density of the aspartate and the typical kink of the proline that precedes the motif B (the aspartate of motif B is S or V in the other motifs, the proline is a Q in motif D). Motif A (^664^LIDL^667^) directly precedes motif B, and Leu664 and Ile665 of motif A of the GTSE1 peptides of monomers C and D are actually occupying Site 4 of a symmetry related monomer, where Leu664 sits in the space of Phe8 of an amphiphysin peptide when superimposed with the 5M5T structure (sequence ^3^LLDLD**F**LE^10^). This crystal packing effect leads to a most likely artificial conformation of motif A of the GTSE1 peptide. In contrast, Site 4 of monomers C and D is empty, indicating a low intrinsic affinity of the GTSE1 peptide to this region. Similarly, Site 2 at the center of the beta-propeller of clathrin is either occupied by a symmetry-related molecule or empty, except for some very fragmented and weak density.

For monomers A and B, the density in site 1 most likely corresponds to motif C (^686^SRP**LIDL**M^693^), based on the shorter side chain density for the Phe/Leu position of motifs B/C and the additional density for the methionine 693 side chain. In this case, there is no additional density visible for the residues preceding the LIDL motif. A possible explanation for the presence of motif C in site 1 of two of the clathrin monomers could be that in the crystal the C-terminus of the motifs A/B (cysteine 673) that is bound to site 1 of monomer C is quite close (12Å) to the N-terminus of the motif C peptide that is bound to site 1 of monomer A, and, similarly, a GTSE1-peptide comprising motifs A to C could easily bridge the binding sites 1 on monomers B and a symmetry-related monomer C. Thus, motif C might be preferred in site 1 due to the interaction of the neighboring motifs A/B with another site 1 on a neighboring molecule in the crystal. This corresponds to the observation that the GTSE1-C2 construct (motif C alone) is not able to pull down clathrin (Fig 2C).

In summary, it seems that motif D of GTSE1 is always preferred in site 3, but that site 1 is more promiscuous and can bind either motif B or motif C, depending on the environment in the crystal. It could still be possible that motif A confers additional affinity of the GTSE1 peptide to clathrin in solution since the Asp666 of motif A is close to Lys98 that is part of a basic patch of clathrin (Fig Struct S3). The position of the LIDF/L motifs in site 1 and 3 are very similar to a clathrin-amphiphysin peptide complex structure (sequence ^3^LLDLD**F**LE^10^, PDB ID 5M5T), but, in contrast to the amphiphysin peptide, the GTSE1-peptide in site 3 has an extended contact area with clathrin due to the additional N-terminal residues “PVV” that are clearly visible in the electron density. For GTSE1, it is possible that the two peptides in site 1 and site 3 could belong to the same GTSE1 molecule since the distance of approximately 50 Å can be easily bridged by the 32 amino acids between motifs B and D. If motif C would be preferred in site 1, a simultaneous binding of motifs C and D to site 1 and site 3 of the same clathrin molecule would not be possible, suggesting that this configuration in the crystal might be an artifact, as explained above. Since in solution the intramolecular binding should be strongly preferred over formation of intermolecular complexes, one would expect the dominant species to be motif D in site 3 and motifs A/B in site 1.

**Fig Struct S1:**
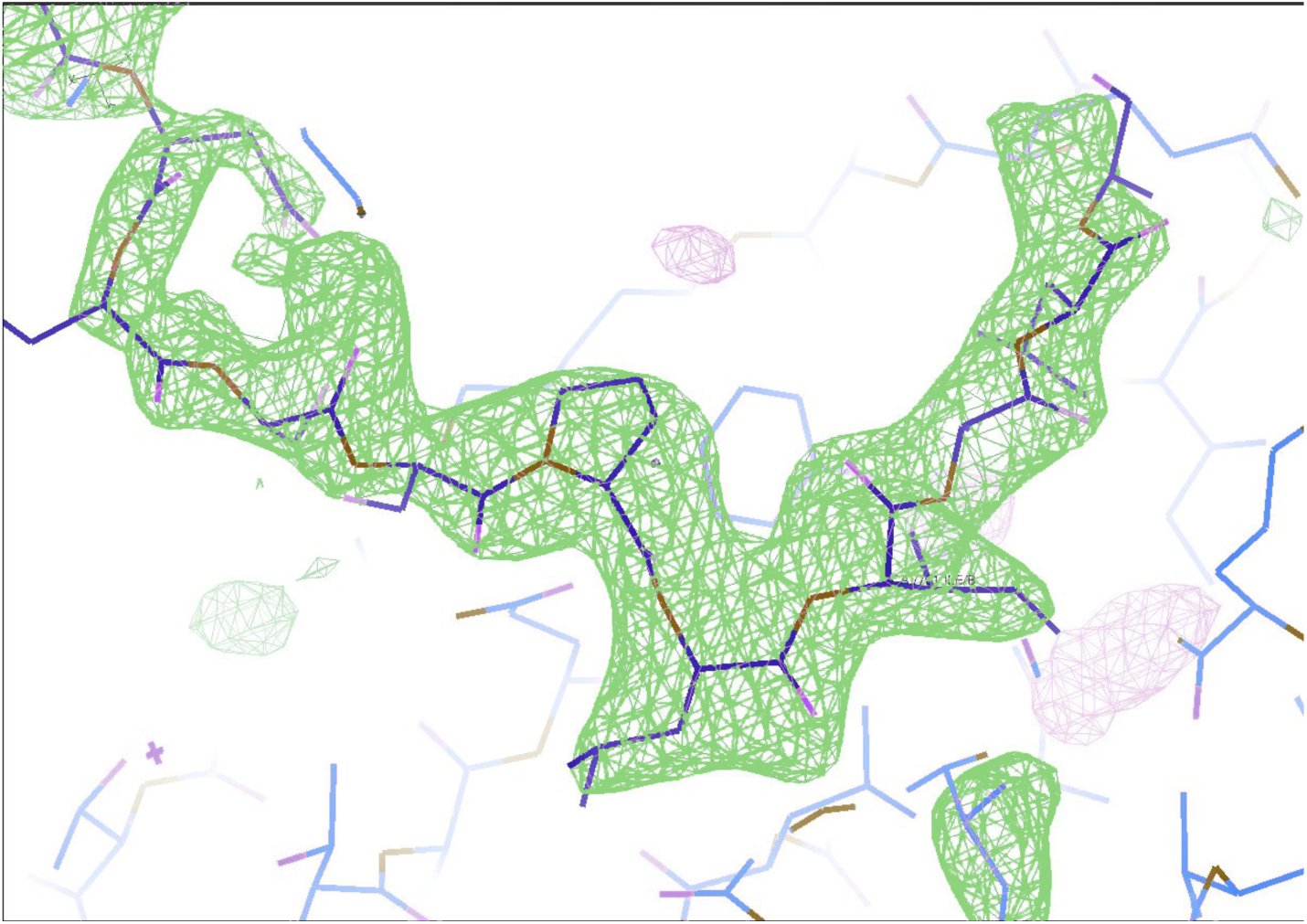
Fo-Fc density for the GTSE1-C4-A-E peptide.

**Fig Struct S2:**
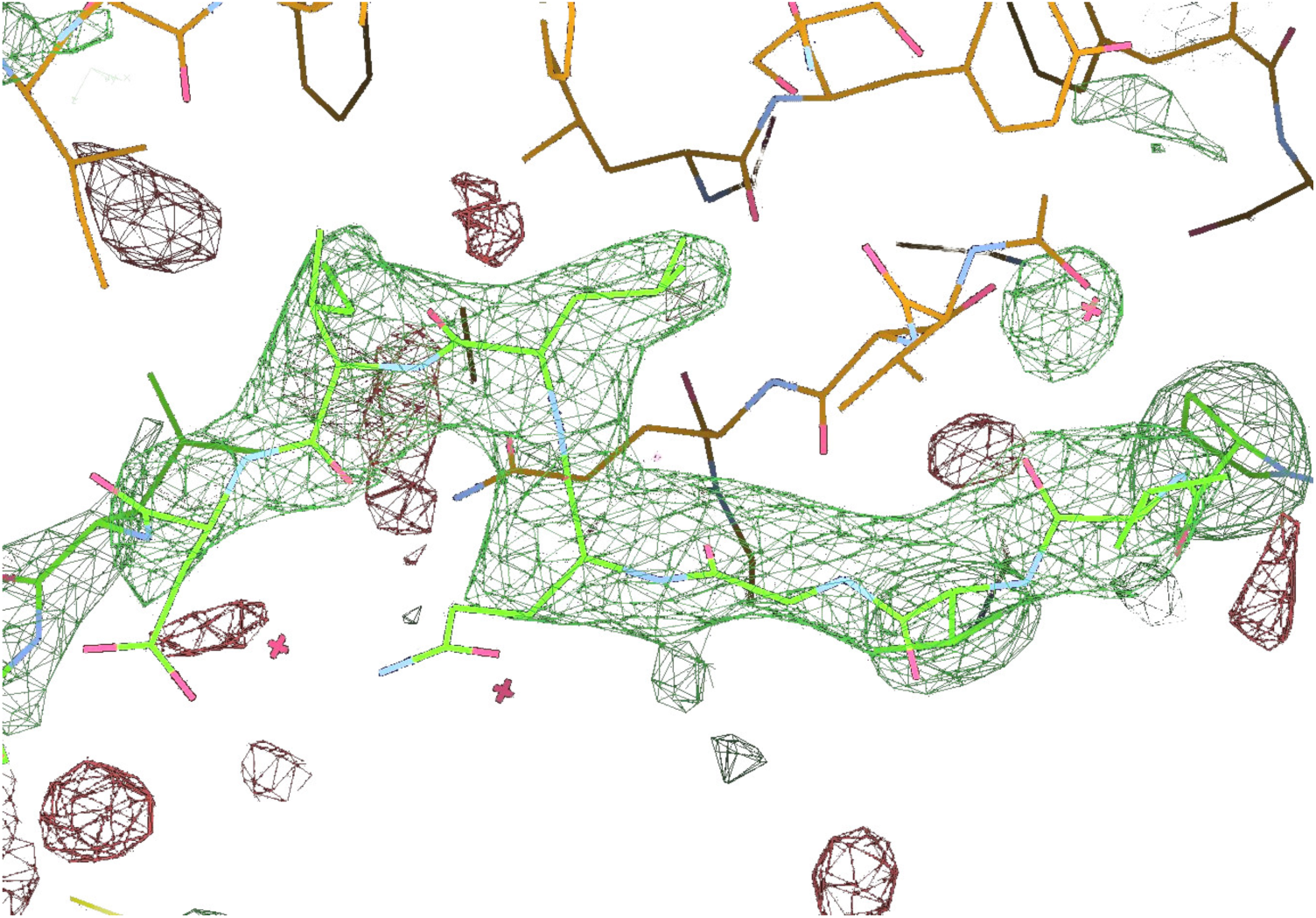
Fo-Fc density for the GTSE1-C5-A-D peptide: monomer A – motif D in site 3.

**Fig Struct S3.**
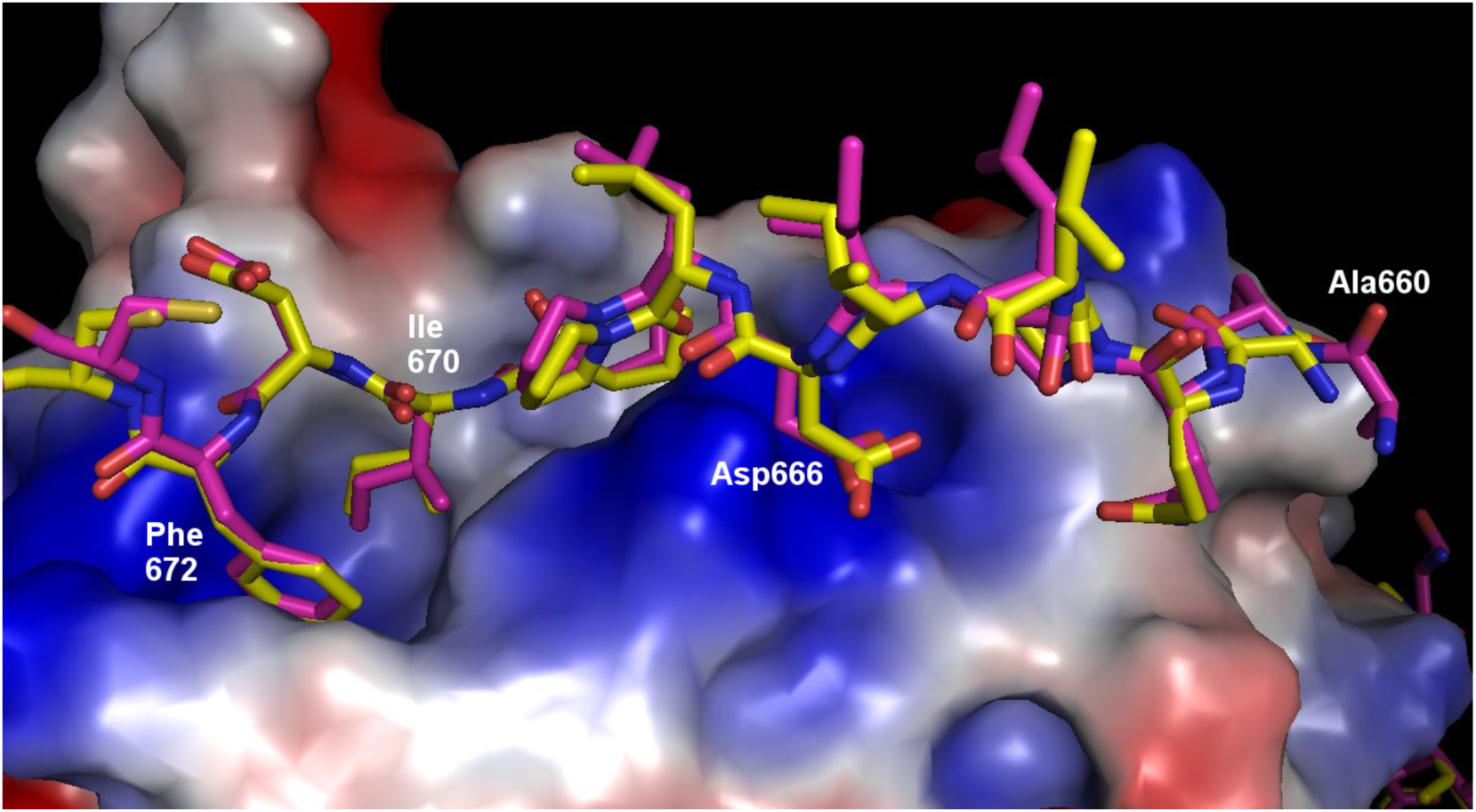
GTSE1-C5 motif A (^664^LIDL^667^) and motif B (^669^LIDF^672^), with Asp666 of motif A interacting with a basic patch on clathrin (monomer D, site 1). Figure created with PyMOL (Schrödinger, LLC).

